# Model selection reveals selective regulation of blood amino acid and lipid metabolism by insulin in humans

**DOI:** 10.1101/2023.09.18.558212

**Authors:** Suguru Fujita, Ken-ichi Hironaka, Yasuaki Karasawa, Shinya Kuroda

**Affiliations:** Department of Biological Sciences, Graduate School of Science, The University of Tokyo, Tokyo, 113-0033, Japan; Department of Biotechnology, Graduate School of Agricultural and Life Sciences, The University of Tokyo, Tokyo, 113-8657, Japan; Department of Neurosurgery, Graduate School of Medicine, The University of Tokyo, Tokyo, 113-0033, Japan

## Abstract

Insulin plays a crucial role in regulating the metabolism of blood glucose, amino acids, and lipids in humans. However, the mechanisms by which insulin selectively controls these different metabolites are not yet fully understood. To address this question, we used mathematical modeling to identify the selective regulatory mechanisms of insulin on blood amino acids and lipids. Our study revealed that insulin negatively regulates the influx and positively regulates the efflux of lipids, which is consistent with previous findings. In contrast, we did not observe the previously reported negative regulation of BCAA influx by insulin; instead, we found that insulin positively regulates BCAA efflux. We also observed that the earlier peak time of lipids compared to BCAAs is dependent on insulin’s negative regulation of their influx. Overall, our findings shed new light on how insulin selectively regulates the levels of different metabolites in human blood, providing valuable insights into the pathogenesis of metabolic disorders and potential therapeutic interventions.

## INTRODUCTION

Insulin is a hormone that plays a key role in regulating the body’s metabolism^1,2^. One of insulin’s primary functions is to regulate blood glucose levels, but it also has a role in regulating other blood metabolites such as amino acids and lipids^3–6^. The previous studies have shown that the regulation of these metabolites by insulin is selective and can vary over time after oral glucose ingestion^7^. While there has been extensive research on the mathematical modeling of insulin’s regulation of blood glucose^8–13^, there has been limited investigation of its selective regulation of blood amino acids and lipids^14–17^. To address this gap, we utilized mathematical model selection to explore insulin’s selective regulatory mechanisms on blood amino acids and lipids, considering their temporal patterns after oral glucose ingestion.

Numerous studies have investigated how insulin regulates amino acids and lipids in the bloodstream^4,18,19^. Insulin reduces blood amino acid concentrations by limiting the release of amino acids into the bloodstream from skeletal muscle^18,20^ and promoting protein synthesis in the liver and other tissues^3^. Specifically, insulin inhibits the release of leucine, isoleucine, methionine, tyrosine, phenylalanine, and threonine^18,20^ from skeletal muscle, while promoting protein synthesis from amino acids in the liver and other tissues^3,4,7,18^. In addition, insulin’s inhibitory effects on glycogenesis and urea synthesis have been shown to reduce the concentrations of arginine, citrulline, and ornithine in the blood^4,21^. These findings have been demonstrated through the use of various methods, including oral glucose ingestion studies^3,4,7,18,19,21^. In the same metabolic group, leucine and isoleucine showed similar temporal patterns, while ornithine and citrulline showed different temporal patterns from leucine and isoleucine after glucose ingestion^7^. These differences in metabolic regulatory mechanisms of amino acids are reflected in their temporal patterns after glucose ingestion^7^.

Blood lipids consist of free fatty acids and ketone bodies, such as 3-hydroxybutyrate. Insulin plays a crucial role in regulating blood lipid metabolism. With regards to free fatty acids (FFA), insulin reduces their concentration in the blood by inhibiting their efflux from adipose tissue into the blood and by promoting their accumulation as triglycerides (TAG) in adipose tissue^5,6^. During feeding, insulin inhibits the activity of hormone-sensitive lipases that regulate lipolysis in adipose tissue and also inhibits the synthesis and release of FFA into the blood by degrading TAG^22^. In addition, insulin indirectly regulates FFA synthesis from TAG by regulating blood glucose levels^5^. For ketone bodies, insulin decreases their concentration in the blood by inhibiting ketogenesis in the liver^23–25^. Insulin also promotes the utilization of ketones, increasing their removal rate from the blood^26^. Taken together, amino acids and lipids such as FFA and ketone bodies showed different temporal patterns, while lipids showed the similar temporal patterns to each other after glucose ingestion^7^. These differences in metabolic regulatory mechanisms of lipids and amino acids are reflected in their temporal patterns after glucose ingestion^7^.

Mathematical models can help identify the regulatory mechanisms of blood metabolites by insulin. By analyzing the temporal patterns of blood metabolites, mathematical models can estimate the model structure and parameters, allowing researchers to infer the selective regulatory mechanisms of insulin. While there have been studies using mathematical models for the metabolic regulation of insulin and glucose^8–13^, there has been less research on the regulation of blood amino acids^14^ and lipids^15–17^ by insulin. Several mathematical models have been developed for the kinetics of blood amino acids and lipids, but none have explained the differences in the temporal patterns of these metabolites after oral glucose ingestion. One study estimated a phenomenological regulatory structure for blood amino acids without considering metabolic map information^27^, but a comprehensive analysis of the selective regulation of amino acids and lipids by insulin using mathematical model selection and detailed blood metabolite data has not yet been performed.

In this study, we used time course data of blood metabolites and hormones from three healthy human subjects who ingested 3 doses of glucose with rapid or slow ingestion to identify the regulatory mechanisms of blood metabolites by insulin^28,29^. We used mathematical model selection to compare different regulatory models based on the metabolic map and statistically selected the best model. We found that BCAAs are positively regulated in terms of efflux, while lipids are positively regulated in terms of efflux but negatively regulated in terms of influx. This regulation pattern for BCAAs is consistent with previous studies^3^, while the negative regulation of insulin reported in a previous study^4^ was not necessary to explain the influx of amino acids in this study’s dataset. This suggests that insulin effectively stimulates the efflux of BCAAs rather than inhibiting their influx. The regulation of lipid, citrulline, and methionine in the selected model is also consistent with previous studies^3^. By using mathematical model selection and glucose dose-dependent time course data of blood metabolites, we were able to infer the effective mechanisms of selective metabolic regulation.

## RESULTS

### Blood metabolites data

In order to perform model selection, we used a dataset from our previous study which included time course data of blood hormones and metabolites in three healthy human subjects^7,29,30^. The dataset included 14 amino acids such as leucine and valine, and 4 lipids including FFA and ketone bodies, which have distinct time patterns based on previous studies (Fig. 1, Fig. S1a, see Methods). The dataset included 14 amino acids such as leucine and valine, and 4 lipids including FFA and ketone bodies, which have distinct temporal patterns based on previous studies (Fig. 1a, Fig. S2)^7,29^. Lipids were found to peak earlier and return to fasting values faster than amino acids (Fig. 1a). For instance, in the case of 75 g Bolus, amino acids peaked later than 120 minutes and did not return to fasting values within 240 minutes, while lipids peaked earlier than 120 minutes and returned to fasting values within 240 minutes (Fig. 1a, arrowhead). These temporal differences suggest that there are selective regulatory mechanisms for amino acids and lipids. Using this dataset, we performed model selection to explain the regulatory mechanisms by analyzing the time series data.

**Fig. 1.**
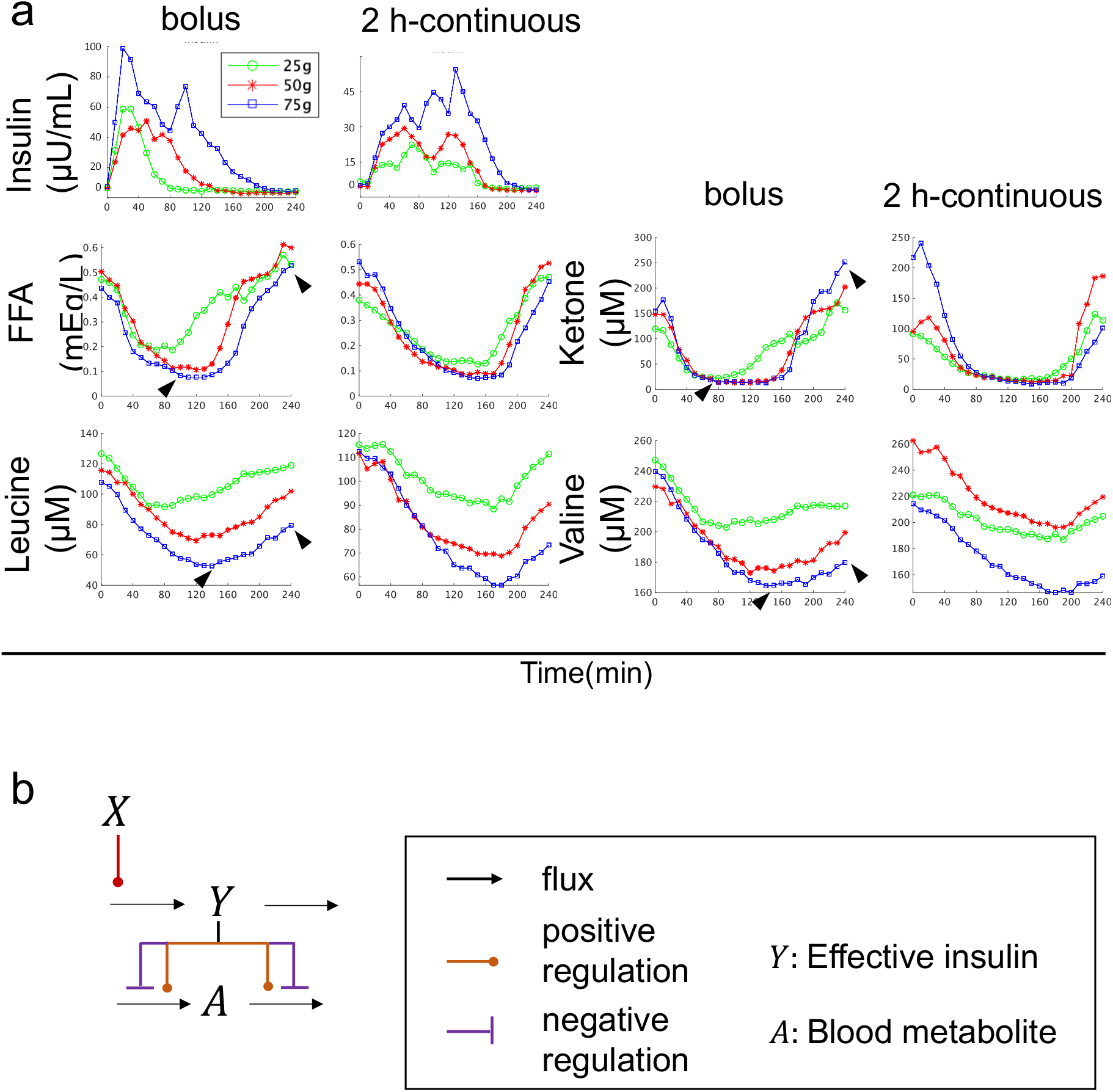
Blood metabolites data and mathematical model structure for model selection. **a** Time course data on the mean values of blood insulin and blood metabolites by glucose ingestion^29^. The doses and ingestion patterns are indicated at the top. Green, red, and blue indicate three different doses: 25 g, 50 g, and 75 g, respectively. See text for arrowheads. **b** Model structure for model selection. (see Methods).

### Mathematical model structure for model selection

In this study, we developed a mathematical model using ordinary differential equations (Fig. 1b, Fig. S3, see Methods) represented by the S-system, which is a type of power-law formalism^31,32^. As there could be multiple mechanisms of insulin action on blood amino acids and lipids, we developed several alternative models (Fig. S3), each including blood insulin (X), effective insulin (Y), and blood metabolite (A), (Fig. S3). These models differed in whether effective insulin regulated the influx or efflux of the blood metabolite (Fig.S3, see Methods). We used time series data of blood insulin as input (Fig. 1, Fig. S3, see Methods) and constructed a regulatory model for each blood metabolite that best fit the average temporal pattern of each metabolite across three healthy human subjects. We estimated the parameters of each model separately to fit the time course data of each metabolite (Fig. 1, S4). The best model was selected by minimizing the Akaike Information Criterion (AIC), which takes into account the complexity of the model and its fit to the time course of each metabolite (see Methods).

### Selected model of each blood metabolite

The best models for the blood metabolites were sorted into five groups based on their regulation by insulin (Fig. 2, Table S1). In the models of ornithine and tyrosine (model #1), both the influx and the efflux were positively regulated by effective insulin (Y) (Fig. 2, Fig. S5). In the lipid model (model #3), the influx was negatively regulated and the efflux were positively regulated by effective insulin (Fig. 2, Fig. S5), while the models for arginine and proline (model #4) showed that both the influx and efflux were negatively regulated by effective insulin(Fig. 2, Fig. S5). The models for BCAAs (model #5) showed that the influx was not regulated by effective insulin, but the efflux was positively regulated (Fig. 2, Fig. S5). In the models of amino acids such as serine and threonine (model #8), the influx was negatively regulated while the efflux was not regulated by effective insulin. Taken together, BCAAs have positive regulation of the efflux, while lipids have positive regulation of the efflux and negative regulation of the influx. We compared our selected model structures with previous knowledge (Fig. S5). For lipid, negative regulation of the influx and positive regulation of the efflux has been reported^5,22–26^, consistent with our result of the selected model. For citrulline and methionine, negative regulation of the influx has been reported^4^, consistent with our result (Fig. S5). For BCAAs, positive regulation of the efflux has been reported^3^, consistent with our result. However, the negative regulation of influx by insulin has been reported^4^, whereas it was dispensable in the model (Fig. S5). Our results suggest that insulin may only effectively stimulate the efflux of BCAAs, rather than inhibit the influx.

**Fig. 2.**
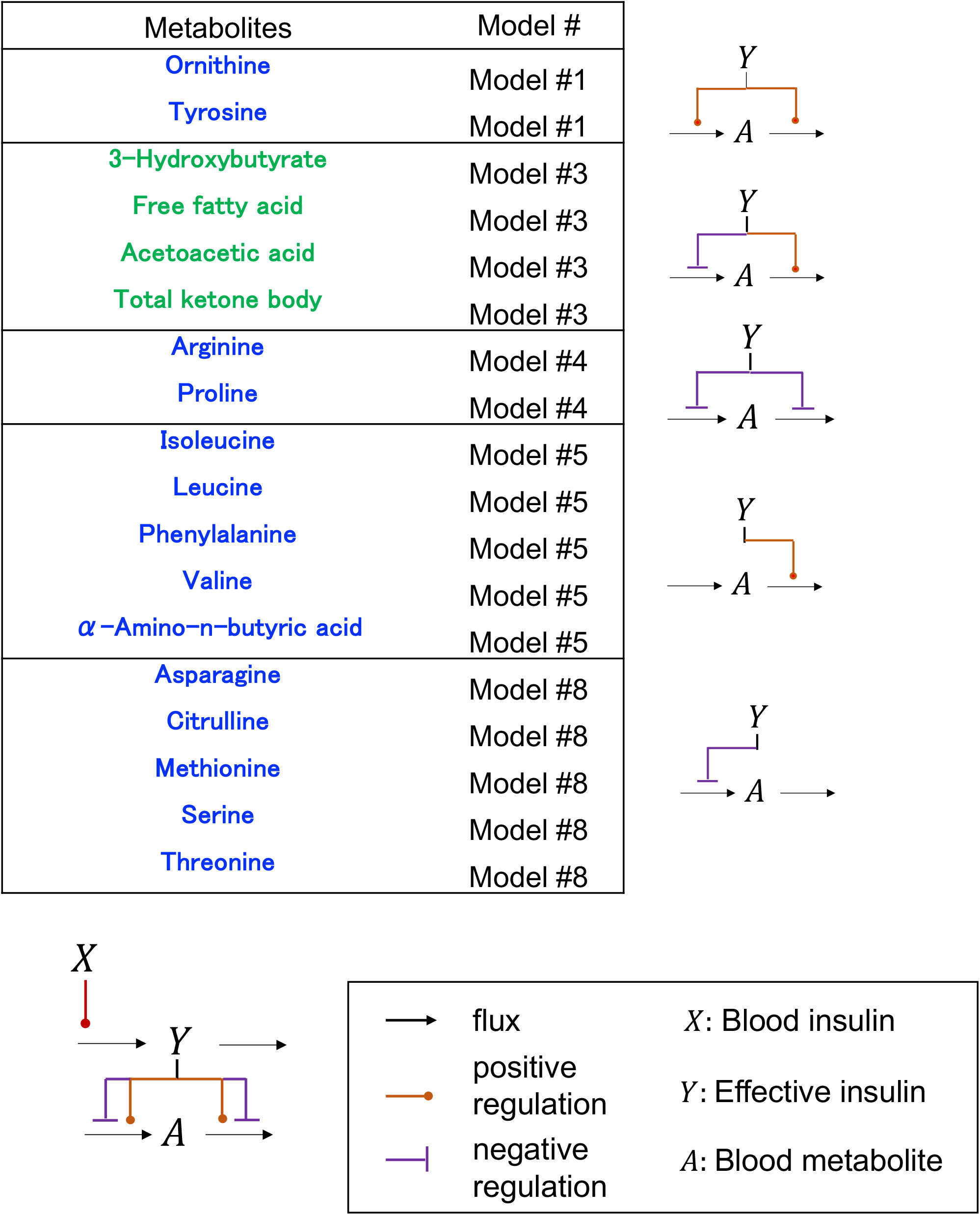
The model structures of the selected model. The model structure with each number models are shown in (Fig. S4).

### Model parameters reflecting the features of temporal pattern

We analyzed the relationship between the estimated model parameters and experimental features extracted from our previous study using tensor decomposition (Fig. 3, Fig. S1b)^29^. The features, called ‘Feature 1’ and ‘Feature 2’, correspond to *y*_*l*_4_*m*_, *l*_4_ = 1,2, respectively (Eq.1 in Fig. S1b)^29^. Since Feature 1 represents the most dominant feature of the dataset, we focused on Feature1 in this study. Feature 1 reflects the peak time of temporal patterns of blood metabolites, with higher Feature 1 values indicating later peaks of similar temporal patterns among individuals and experimental conditions (Fig. S1b).

**Fig. 3.**
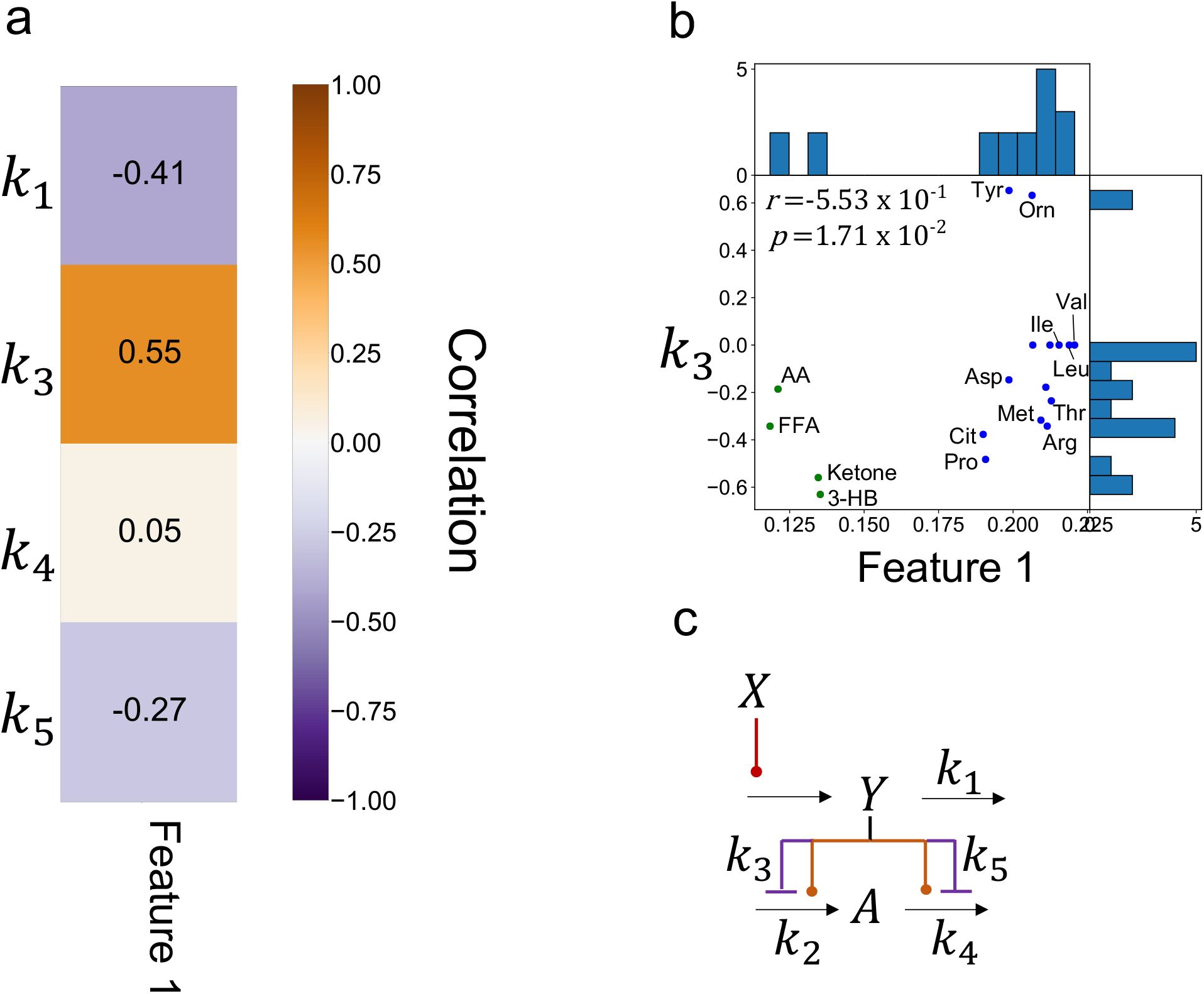
Relationship between model parameters and physiological features. **a** Heat map of correlation coefficients between model parameters and features. **b** Scatter plot of *k*_3_ with Features 1 of the indicated metabolites. *r* and *p* indicate the correlation coefficient and *p* value, respectively. Abbreviations for the representative molecules as follows: Asp aspartic acid; Cit, citrulline; FFA, free fatty acid; 3-OH, 3-hydroxybutyric acid; Ketone, Total ketone body; Glu, glutamic acid; His, histidine; Ile, isoleucine; Ins, insulin; Leu, leucine; Tyr, tyrosine; Val, valine. The dot’s colors correspond to the metabolic group (blue: amino acids, green: lipids). **c** Model diagram with model parameters.

One of the model parameters, *k*_3_, was found to be strongly correlated with Feature 1, which are experimental features reflecting the peak time of temporal patterns of blood metabolites (Fig. 3, correlation coefficients *r* = 0.55 for Feature1, *P* < 0.05). Specifically, a greater negative strength of regulation on the influx of effective insulin (represented by a larger *k*_3_ value) was associated with earlier peaks in both similar temporal patterns among experimental conditions. This is because insulin transiently increases after glucose ingestion (Fig. 1), and the greater the strength of regulation of effective insulin on the influx, the more transient (*i*.*e*., earlier peak) the downstream metabolites’ temporal patterns (Fig. 1, Fig. S1b). Lipids with a negative *k*_3_ parameter, indicating a large strength of regulation of effective insulin on the influx, showed earlier peaks than BCAAs with a *k*_3_ parameter of 0, indicating no strength of regulation of effective insulin on the influx. These results suggest that the regulation of effective insulin on the influx contributes to earlier peaks in lipid temporal patterns compared to BCAA temporal patterns (Fig. S1b).

### The regulation of the amplitude of amino acids and lipids by amplitude of insulin

We have previously found that the time course of different group of blood metabolites in our study is characterized by “amplitude” and “rate”^7^. Therefore, we examined which regulation by insulin in the model is responsible for amplitude and rate of each metabolites (Figs. 4 and 5). We used the mathematical model (Fig. 4) to investigate how insulin regulates different temporal patterns of metabolites. We first looked at how the amplitude of insulin affects the amplitude of metabolites. The amplitude of metabolites was quantified for various amplitudes of insulin (Fig. 4b,c)^33^. The dose-response curves for all metabolites showed a monotonic increase as the amplitude of insulin increased (Fig. 4d, Fig. S7b). We calculated the EC_50_, which represents the half-maximal concentration of insulin required for half maximum amplitude of the metabolites (Fig. 4e, Fig. S7c, see Methods). It should be noted that we normalized the amplitude of the metabolites corresponding to the maximum amplitude of insulin to 1.

**Fig. 4.**
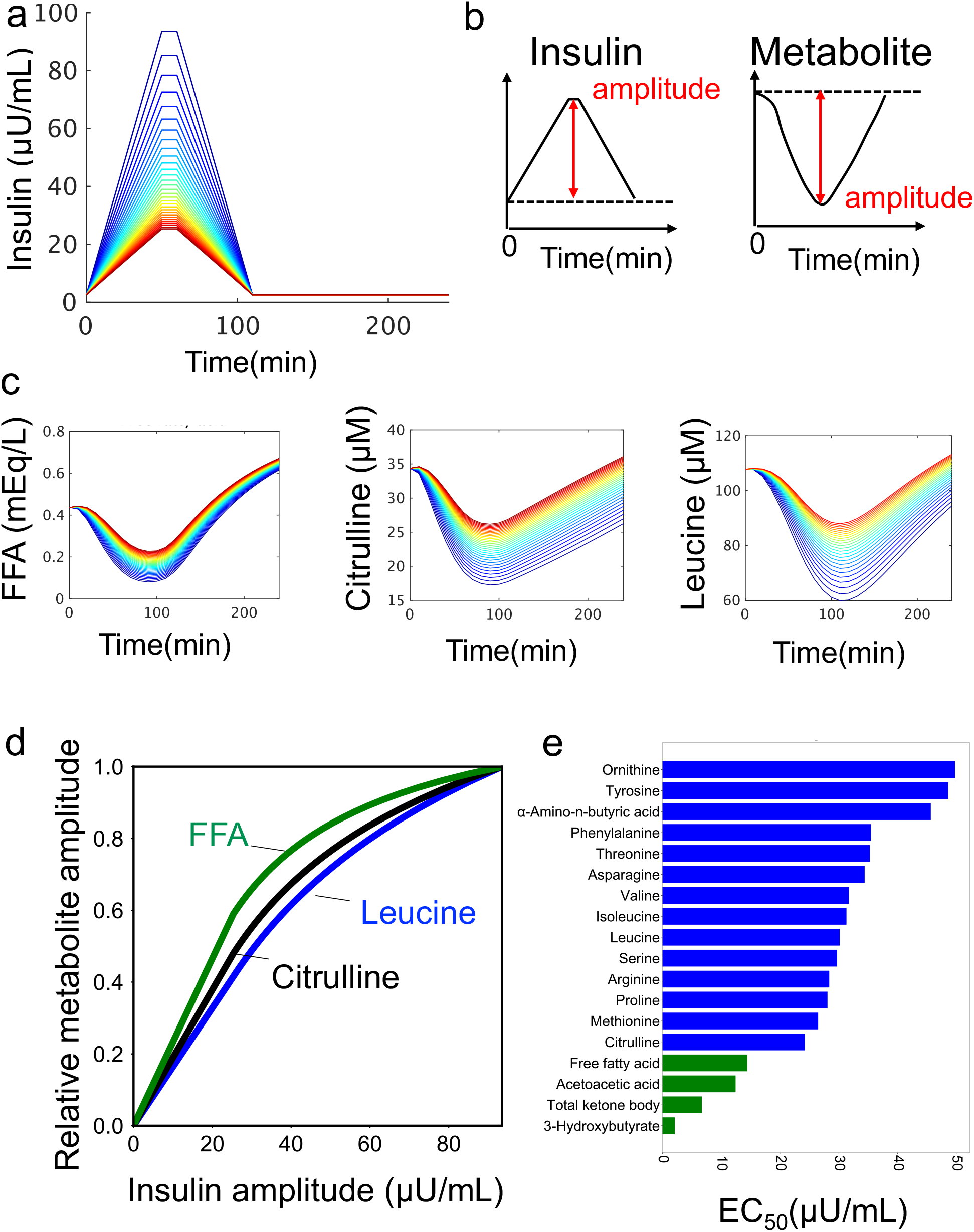
The regulation of the amplitude of the metabolites by amplitude of insulin. **a** Temporal pattern of insulin with different amplitudes as input. The colors of lines indicate different peaks. **b** Definition of amplitudes in time series for insulin and metabolites **c** Temporal pattern of metabolites as outputs. The colors of lines correspond to the colors of the inputs(**a**). **d** The amplitudes of the indicated metabolite against the amplitudes of insulin in the simulation. **e** EC_50_s of metabolites against the amplitude of insulin. The color of the bar indicates metabolic group (blue: amino acids, green: lipids)

According to the results of our investigation using the mathematical model (Fig. 4), we found that the EC_50_ values for amino acids and lipids were 34.23 (μU/mL) and 8.92 (μU/mL), respectively (Fig. 4e). Amino acids had larger EC_50_ values compared to lipids, which means that they can respond to a wider range of insulin’s amplitudes than lipids (Fig. 4e). Here we calculated average values for each metabolic group. Conversely, lipids had smaller EC_50_ values, indicating that they are more sensitive to the lower amplitude of insulin (Fig. 4e). Taken together, our findings suggest that insulin can selectively regulate amino acids and lipids based on the amplitude of insulin, and citrulline shows intermediate characteristics between amino acids and lipids (Fig. 4d,e).

### The regulation of the rates of amino acids and lipids by the increasing rate of insulin

In the next step, we examined how the increasing rate of insulin is linked to the rate of metabolites (Fig. 5 a,b). To quantify the rate of metabolites, we introduced an index called ‘RI (Rate Index),’ which represents the time duration required for a metabolite to transition from 25% to 75% of its maximum response (Fig. 5c)^33^. We investigated how the rate of metabolites is affected by the increasing rate of insulin. The duration time of insulin was used as a measure of its increasing rate, with shorter durations indicating faster increasing rates (Fig. 5a,c). The results showed that as the duration time of insulin increased (*i*.*e*., the increasing rate of insulin decreased), the RIs of metabolites also decreased, indicating that the rate of metabolites is influenced by the increasing rate of insulin (Fig. 5d, S7d). The dynamic range of RI was then calculated as a measure of how much information regarding the rate of increase of insulin is transferred to metabolites (Fig. 5e). Lipids had a larger RI and a larger dynamic range than amino acids, suggesting that insulin can more finely control the rate of amino acids than lipids by changing the increasing rate of insulin (Fig. 5d,e).

**Fig. 5.**
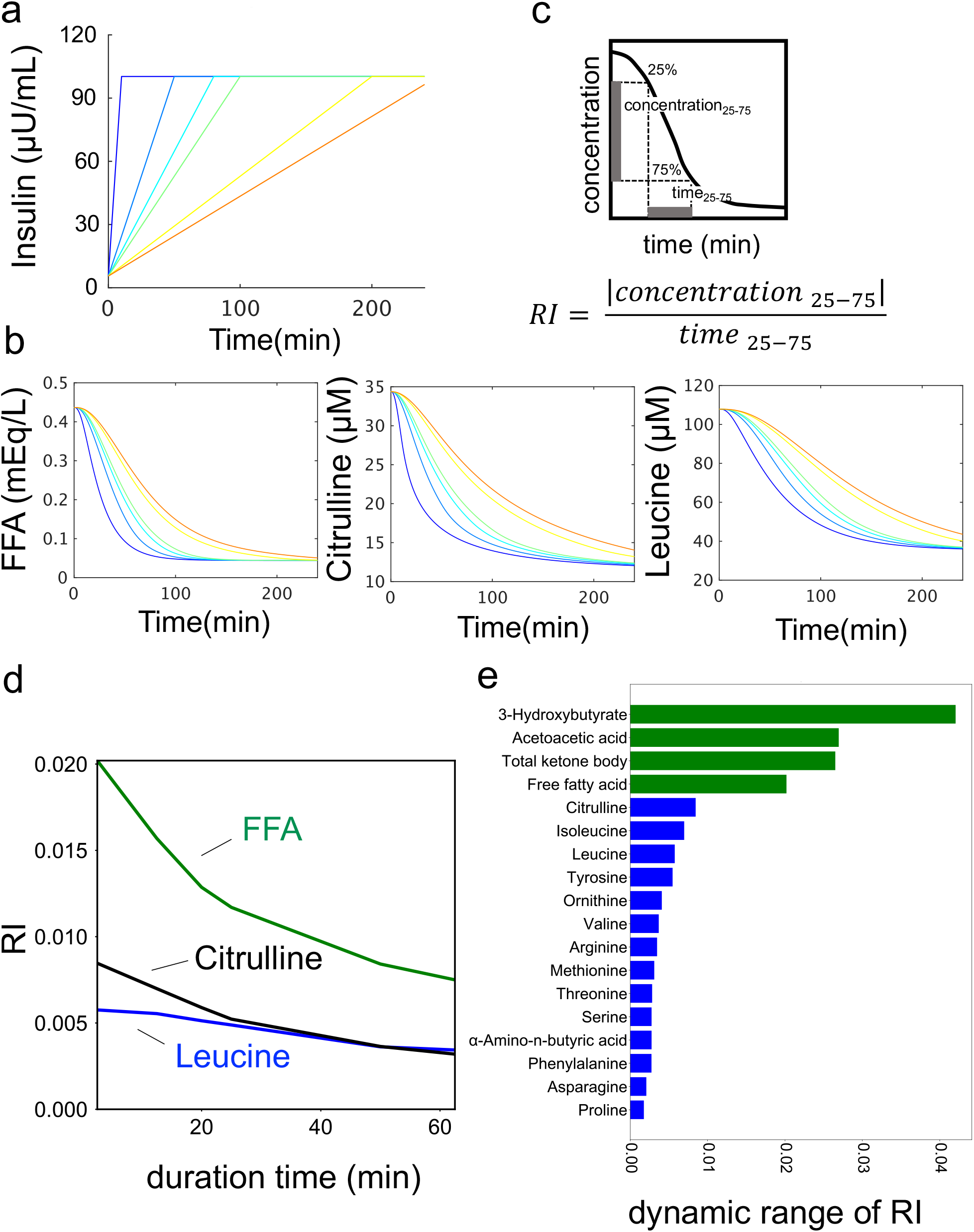
The regulation of the rates of amino acids and lipids by the increasing rate of insulin. **a** Temporal patterns of insulin with different increasing rates as input. The colors of lines indicate different rates. **b** The definition of Rate Index (RI) in time series for insulin and metabolites **c** Temporal pattern of metabolites as outputs. The colors of lines correspond to the colors of the inputs(b). **d** The RI of the indicated metabolite against the rate of insulin in the simulation. The x-axis of shows the duration of insulin, which is inversely proportional to the rate of insulin increase. **e** Dynamic range of RI of metabolites. The color of the bar indicates metabolic group (blue: amino acids, green: lipids)

### Model validation

We examined the validity of the selected model for each metabolite by applying to different datasets (Fig. 6) from five healthy individuals who either ingested 75g of glucose rapidly or over 2 hours, which are not used for model training (Fig. S8, see Methods). The selected model successfully reproduced the peaks at around 60 minutes for lipids such as FFA and ketones, and at 150 minutes for amino acids such as leucine and isoleucine for the bolus condition (Fig. 6, arrowhead). For the 2-hour continuous condition, the model reproduced the peak around 120 minutes for lipids and the peak around 180 minutes for amino acids, further confirming the validity of the model (Fig. 6, arrowhead).We compared the experimental and simulated values using the selected model for each metabolite and found a high correlation (*r* > 0.9) for all metabolites (Fig. S9, S10), providing further evidence for the accuracy of the selected model.

**Fig. 6.**
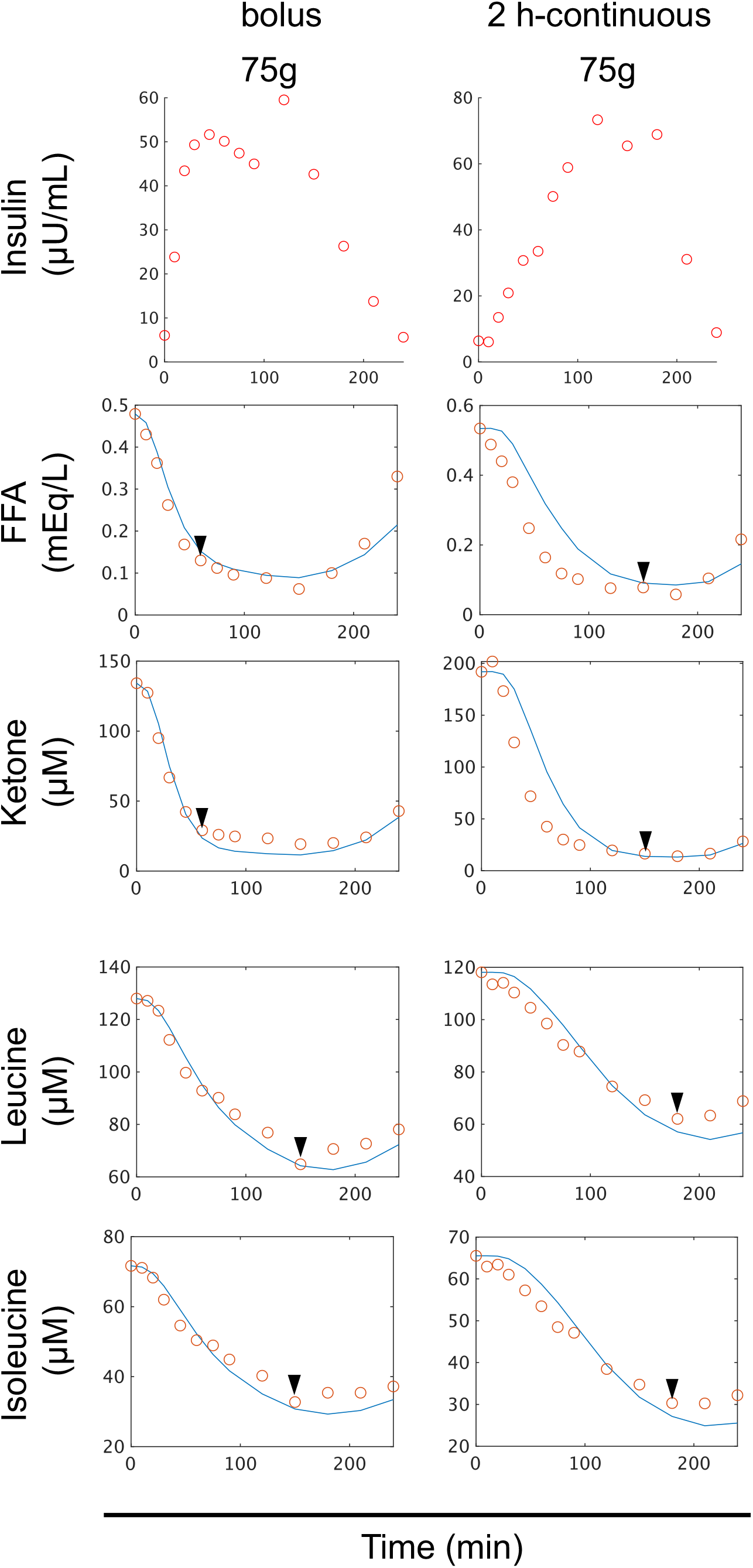
Model validation. Time course data on the mean values of blood insulin and blood metabolites in five individuals by glucose ingestion for validation. The doses and ingestion patterns are indicated at the top. The blue lines indicate the temporal patterns of simulations, and the red circles indicate the time course data of experiments. Before and after glucose ingestion, the concentration of blood insulin and 14 amino acids, including leucine and valine, and 4 lipids, including FFA and ketone bodies were measured at 14 time points from 10 min before fasting to 240 min after glucose ingestion (-10, 0, 10, 20, 30, 45, 60, 75, 90, 120, 150, 180, 210, 240 minutes).

## DISCUSSION

In this study, we developed a mathematical model to analyze changes in blood metabolites over time using data from three healthy human individuals who consumed three different doses of glucose at varying rates (Fig. 1, Fig. S4). The selected model structures varied between groups of blood metabolites, indicating that insulin selectively regulates different groups of metabolites (Fig. 2). Interestingly, the same model structure was chosen for amino acids such as BCAA, as well as for lipids such as FFA and ketone bodies, suggesting that while the regulation of insulin is different between metabolic groups such as amino acids and lipids, it is similar within each metabolic group (Fig. 2).

In this study, we analyzed the correlation between estimated model parameters and features extracted by tensor decomposition (Fig. 3, Fig. S1)^29^. We found that the model parameter, *k*_3_, was highly correlated with Feature 1. *k*_3_ represents the strength of the negative regulation of effective insulin on the influx of metabolites. This indicates that the differences in the temporal patterns of amino acids and lipids (Fig. 2, Fig. S1b) can be explained by variations in the strength of the regulation of effective insulin on the influx. These findings demonstrate how data-driven features obtained by tensor decomposition can be used to drive hypotheses and uncover underlying physiological mechanisms.

Our study focused on examining the role of insulin in regulating the influx and efflux of blood metabolites in a mathematical model. Previous research has shown that insulin inhibits proteolysis in skeletal muscle, which leads to a decrease in the release of amino acids into the blood^4^. In the liver, insulin activates S6 kinase through the AKT pathway, which promotes protein synthesis and causes an increase in amino acid usage for protein synthesis^34^. These findings suggest that insulin negatively regulates the influx of amino acids into the blood while positively regulating the efflux (Fig. S5). However, the selected model structures indicated that insulin only positively regulates the efflux of amino acids from the blood and does not regulate the influx (Fig. 2, Fig. S5). This suggests that while insulin regulation of the efflux is physiologically effective, it is not essential for the influx of amino acids to reflect the data set of this study. Further research is needed to better understand the contribution of insulin regulation to the influx of amino acids in the regulation of blood amino acids.

We found that insulin regulates the blood concentration of lipids such as FFA and ketone bodies through the positive regulation of their influx and the negative regulation of their efflux (Fig. 2). This is consistent with previous study indicating that insulin inhibits fatty acid influx from adipose tissue into the blood^5,6^, leading to decreased blood FFA concentrations and increased triacylglycerol accumulation in adipose tissue (Fig. S5)^6^. Additionally, insulin has been shown to inhibit ketone bodies synthesis^25,35^ and increase their removal rate in the blood^26^. Our mathematical model selected the same structures(Fig. 3, Fig. S5), indicating that these regulations of insulin on blood lipids are consistent across multiple studies^5,22–26^.

In this study, we investigated how insulin’s amplitude and rate regulate two components of metabolites (Fig. 4). Differences in the temporal patterns of blood metabolites suggested different regulatory mechanisms by insulin^7^. With respect to the amplitude component, we defined EC_50_ as a measure of sensitivity to insulin and compared it among downstream metabolites (Fig.4, S7). Lipids showed higher sensitivity to insulin than amino acids against the amplitude of insulin, and citrulline showed intermediate sensitivity between lipids and amino acids such as BCAAs (Fig. 4). We used experimental data to demonstrate a similar trend in sensitivity to insulin: that is, the EC_50_ of amino acids is higher than that of lipids (Fig. S7d). As a point of reference, in a previous study in which plasma AA concentrations were measured during normoglycemic insulin infusion in healthy young adult males at four different insulin infusion rates (6, 10, 30, and 400 (mU/(m^2^·min))), the half-maximal response (a value with the same meaning as the EC_50_ in this study) for amino acids was averaged about 30.9 (μU/mL)^18^. The estimates from the model analysis of this study (34.23 (μU/mL)) did not deviate significantly. We also performed a parameter sensitivity analysis of EC_50_ in simulations for each model parameter (Table S2, see Methods). As the parameter sensitivity index, we compared the median value of each parameter for each of the 18 metabolites (Table S2). k_3_ had the highest median value, indicating that the strength of the negative regulation of effective insulin on the influx of metabolites is the most important parameter controlling insulin sensitivity.

The previous studies have shown that insulin’s inhibition of lipolysis and ketogenesis is more sensitive than the inhibition of protein catabolism^25^. We also demonstrated that the sensitivity of citrulline to insulin was intermediate between that of amino acids and lipids^7^. The mathematical model analysis in this study consistently explained the selective regulatory mechanisms for different sensitivities of the metabolites. For the rate component, we defined RI as a measure of the rate component and compared it among downstream metabolites (Fig. 5, S7). Lipids with a larger RI showed a larger dynamic range of increasing rate of insulin than amino acids with a smaller RI. Taken together, our results demonstrate that insulin can more tightly control amino acids than lipids by changing the amplitude of insulin, while insulin can more tightly control lipids than amino acids by changing the increasing rate of insulin.

In our previous studies, we investigated the different temporal patterns of how blood metabolites respond to glucose ingestion using a combination of hypothesis-driven^7^ and data-driven analyses^29^. We discovered that amino acids and lipids selectively decode the amplitude and increasing rate of insulin, respectively. However, we did not fully understand the mechanisms behind this selective decoding. In this study, we used mathematical modeling and time course data of blood metabolites after oral glucose ingestion to provide one explanation for the mechanism of selective regulation of the metabolites by insulin. In this study, we developed a mathematical model based on data from a single dose of glucose (75 g bolus), which is commonly used in clinical settings to assess glucose tolerance. We then compared this model with a model based on multiple doses and durations of glucose (six experiments) data (Fig. S11). The model using multiple doses and durations had a better fit (lower RSS value) for several molecules, including methionine, compared to the model using single dose glucose. This highlights the importance of using multiple data sets to validate mathematical models. The findings indicate that relying solely on data from a single dose of glucose (75g, Bolus) is not enough to accurately capture the dynamics of blood metabolite changes. Instead, data from multiple doses and durations of glucose ingestion are required to better understand the temporal patterns of blood metabolites.

### Limitations of the study

One limitation of this study is the small number of individuals and model structures used. The data were collected from only three individuals due to the time-consuming nature of the experiments, and population-averaged data were used to estimate the model structure without analyzing individual differences. In our previous study^7^, we analyzed individual differences using blood data from 20 individuals. A larger number of individuals would allow for a more comprehensive analysis of individual differences using mathematical models. Additionally, the study assumed only eight simple candidate model structures and did not consider the interaction of insulin with glucose-mediated FFA^36–38^ and mTOR-mediated leucine^39^. While the model successfully reproduced blood amino acid and lipid concentrations, it is possible that the amplitude or rate of insulin altered insulin secretion, clearance, or feedback from FFA or leucine. We investigated the identifiability of the parameters in the model structures selected for each metabolite (Fig. S12)^40,41^. For the lipid model, parameters k_3_ and k_5_ were shown to be identifiable, but k_1_ and k_4_ were found to be indistinguishable; for the amino acid model, including BCAA, parameters k_1_ and k_5_ were shown to be identifiable, but k_4_ was indistinguishable for both valine and leucine. Taken together, the parameters k_3_ and k_5_ representing insulin regulation showed that there are unique solutions in the range tested. We quantified EC_50_ to determine the sensitivity of metabolites to insulin. Due to nonlinear least-squares fitting, lipids showed the smaller EC_50_s than those of the amplitude of insulin, which was given as a simulation input (Fig. 4). In this study, we don’t argue strongly for the abosolute values of EC_50_s of metabolites; however, at least for the fact that lipids have the smaller EC_50_s than amino acids. (Fig. 4, Fig. S7). Further studies are needed to estimate detailed regulatory structures for individual metabolites.

## Supplementary Table legends

**Table S1.**

AIC and RSS for each model of all metabolites

**Table S2.**

Parameter sensitivity analysis for EC_50_.

**Table S3.**

Characteristics of subjects

**Table S4.**

The "data set for model validation"

## Resource availability

### Lead Contact

Further information and requests for resources and reagents should be directed to and will be fulfilled by the Lead Contact, Shinya Kuroda

### Materials availability

This study did not generate new unique reagents.

## EXPERIMENTAL MODEL AND SUBJECT DETAILS

### 1. Subjects

The study involved three healthy subjects for the "data set for model construction" and five healthy subjects for the "data set for model validation". The profiles of the subjects are provided in Table S3. The "data set for model validation" is provided in Tables S4

### 2. Blood sampling

We utilized human blood samples that were obtained from our previous studies^7,29^.

### 3. Ethics committee certification

We followed Japan’s Ethical Guidelines for Epidemiological Research, and the study was approved by the Institutional Review Board and the Ethics Committee of Tokyo University Hospital (Approval No. 10264-(4)). Subjects were recruited through snowball sampling.

## METHOD DETAILS

### 4. Model structure and parameter structure

We developed mathematical models to describe the temporal changes in blood metabolite concentrations (Fig. 1). We developed a total of eight different models, assuming three types of regulation for each blood metabolite’s influx into and efflux from the blood: positive regulation, negative regulation, or no regulation (Fig. S3). We excluded the case where there was no regulation for both influx and efflux. We estimated the model parameters for each model. The specific molecules targeted in our analysis are listed in Fig. 2.

Each regulation type is based on the S-system model^31,32^, and the change in the concentration of a blood metabolite (*A*) is described by the following differential equation:

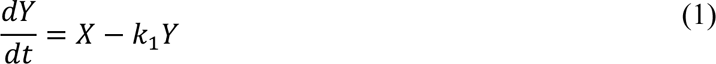

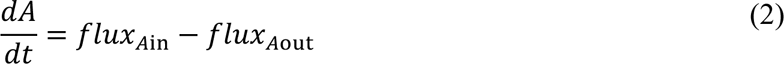

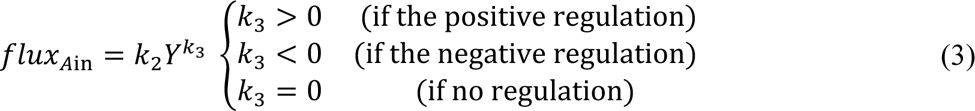

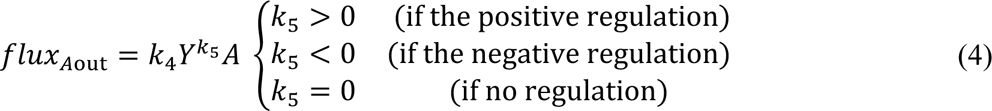

where *X* is blood insulin concentration, *Y* is effective insulin that effectively regulates blood metabolite concentration, and *A* is blood metabolite concentration. The values of *X* and *A* are obtained by averaging the data from three individuals and normalized to have a variance of 1. The term *flux*_*A*in_(*k*_2_*Y*^*k*_3_^) represents the influx of metabolite *A* into the blood and is determined by the positive regulation (*k*_3_ > 0), negative regulation (*k*_3_ < 0), or the reaction rate constant *k*_2_, depending on the effective insulin *Y*. Similarly, the term *flux*_*A*out_ (*k*_4_*Y*^*k*_5_^ *A*) represents the efflux of metabolite *A* from the blood and is determined by the positive regulation (*k*_5_ > 0), negative regulation (*k*_5_ < 0), or the reaction rate constant *k*_4_, depending on the effective insulin Y and blood metabolite concentration *A*.

Using the variables *X*, *Y*, and *A*, and assuming that *X*, *Y*, and *A* are at steady state before glucose ingestion, we determined the initial conditions and parameters based on the estimated parameters and initial values:

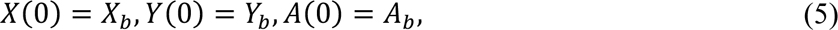

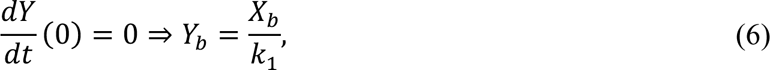

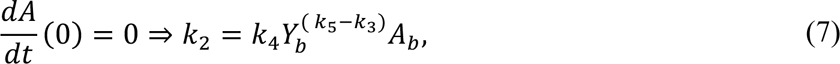

where *X*_*b*_, *Y*_*b*_, and *A*_*b*_ indicate the initial values of *X*, *Y*, and *A*, respectively. Therefore, there are four parameters to be estimated: *k*_1_, *k*_3_, *k*_4_, and *k*_5_. The measured blood insulin concentration was treated as continuous by linear interpolation between the measurement time points, as *X* is used as a continuous value during the simulation, although the actual measured values were used.

### 5. Model selection

We used the Residual Sum of Squares (RSS) as the objective function to minimize the differences between the experimental and simulated values, given by:

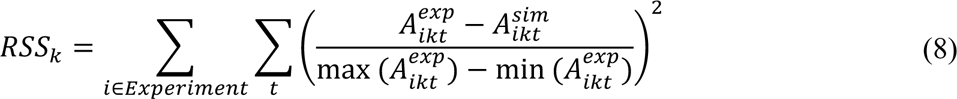

In this equation, 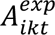 and 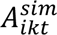 represent the measured and simulated values, respectively, at time *t* and metabolite *k* in experiment *i*, which can take the values 25B, 25C, 50B, 50C, 75B, or 75C. Each experiment is identified by the dose of glucose ingestion and the initial letters of the duration of ingestion (e.g., 25-g-bolus ingestion is denoted as 25B, and 75-g-2-h-continuous is denoted as 75C).

To account for differences in the absolute values of the metabolites across experiments, we normalized the difference between the experimental and simulated values by dividing it by the difference between the maximum and minimum values of the experiment. This normalization helps to mitigate the influence of varying magnitudes in the metabolite concentrations. To find the global optimal solution, we performed parameter estimation using the Evolutionary Programming method^42^. We conducted 20 trials with a parent number of 20 and a generation number of 400. After obtaining the global optimal solution^43^, we further refined it using the simplex search method (Matlab fminsearch) to find a local optimal solution. For each blood metabolite *k*, we performed parameter fitting for all eight models developed in Section 5 using the RSS calculated according to equation (8). To statistically select the regulatory structure of each blood metabolite, we calculated the Akaike Information Criterion (AIC) based on the RSS of each model.

The AIC was computed using the following formula, taking into account that the models being compared have the same total number of data points N for the experimentally measured variables:

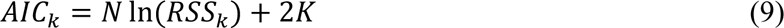

where *K* is the number of parameters in the model that need to be estimated. We assumed that the RSS values follow a normal distribution. The AIC provides a measure of the trade-off between the goodness of fit (represented by the RSS) and the complexity of the model (represented by the number of parameters). By considering both the fit to the data and the model complexity, the AIC allows us to compare and select the most appropriate model for each blood metabolite.

### 6. Parametaer identifiability

The parameter sets that produced the selected models were evaluated for identifiability. We iteratively changed the value of one parameter from its optimal value and re-estimated the remaining parameters^40^. An increase in the cost function (RSS) of the model fit indicates that reliable parameter estimates are obtained and that the parameters are identifiable from the model structure and data.

### 7. Parameter sensitivity analysis

We defined the individual model parameter sensitivity for each subject as follows:

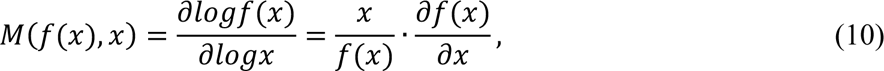

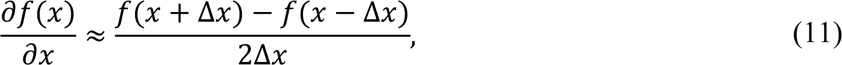

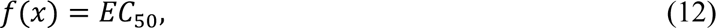

where x is the parameter value and f(x) is EC_50_. The differentiation is numerically approximated by central difference (equation (8), and *x* + Δ*x* and *x* − Δ*x* were set so as to be increased [x (1.1x)] or decreased [x (0.9x)] by 10%, respectively. Finally, we defined the parameter sensitivity by the median of the individual parameter sensitivity for all metabolites. We examined the parameter sensitivity for four parameters. The higher the absolute value of parameter sensitivity, the larger the effect of the parameter on EC_50_.

### 8. The temporal pattern similarity among molecules

In our previous study, we introduced the Temporal Pattern Similarity among molecules (TPSM) as a measure of the similarity between their temporal patterns^7^. The TPSM is defined as follows (Fig. S7a):

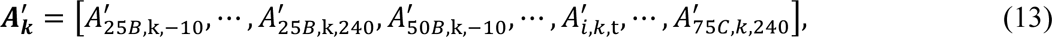

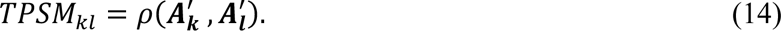

where 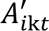 represents the interindividual mean values of the difference from fasting at time *t* for metabolite *k* in experiment *i*. The vector 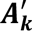 represents the time-series connecting these values (equation 10). The Pearson correlation coefficient, denoted as 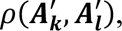 is used to measure the similarity between 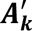 and 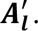 The *TPSM*_*kl*_ (equation 14) represents the temporal pattern similarity between molecule *k* and *l*, indicating how similar their temporal patterns are. Importantly, there were no molecular sets that exhibited a negative correlation.

### 9. Calculation of EC_50_

To determine the half-maximal concentration (EC_50_) of all metabolites in response to insulin, we performed nonlinear least-squares fitting (Fig. S7b,c). It should be noted that although the time series for insulin and metabolites at a peak value of 0 is not depicted in Figure 4, we extrapolated the metabolite peak value to 0 for insulin in order to perform the calculation.

### 10. Data and Code Availability

All data generated or analysed during this study and the code files are included in this article and the article^29^ and their supplementary materials files. The code files used in the simulation are freely available at https://github.com/sfujita0601/ModelSelection_HumanOGTTDose

## ACKNOWLEDGEMENTS

We thank our laboratory members for critical reading of the manuscript. This study was supported by the Japan Society for the Promotion of Science (JSPS) KAKENHI (Grant Numbers JP17H06300, JP17H06299, JP18H03979, JP21H04759), CREST, the Japan Science and Technology Agency (JST) (JPMJCR2123), and by The Uehara Memorial Foundation. S.F receives funding from JST SPRING (JPMJSP2108).

## AUTHOR CONTRIBUTIONS

S.F., K.H. and S.K. analysed the data. S.F., K.H., and S.K. wrote the manuscript. The study was conceived and supervised by S.F., K.H. and S.K.

## DECLARATION OF INTERESTS

The authors declare no competing interests.

## Supplementary Figures

**Fig. S1.**
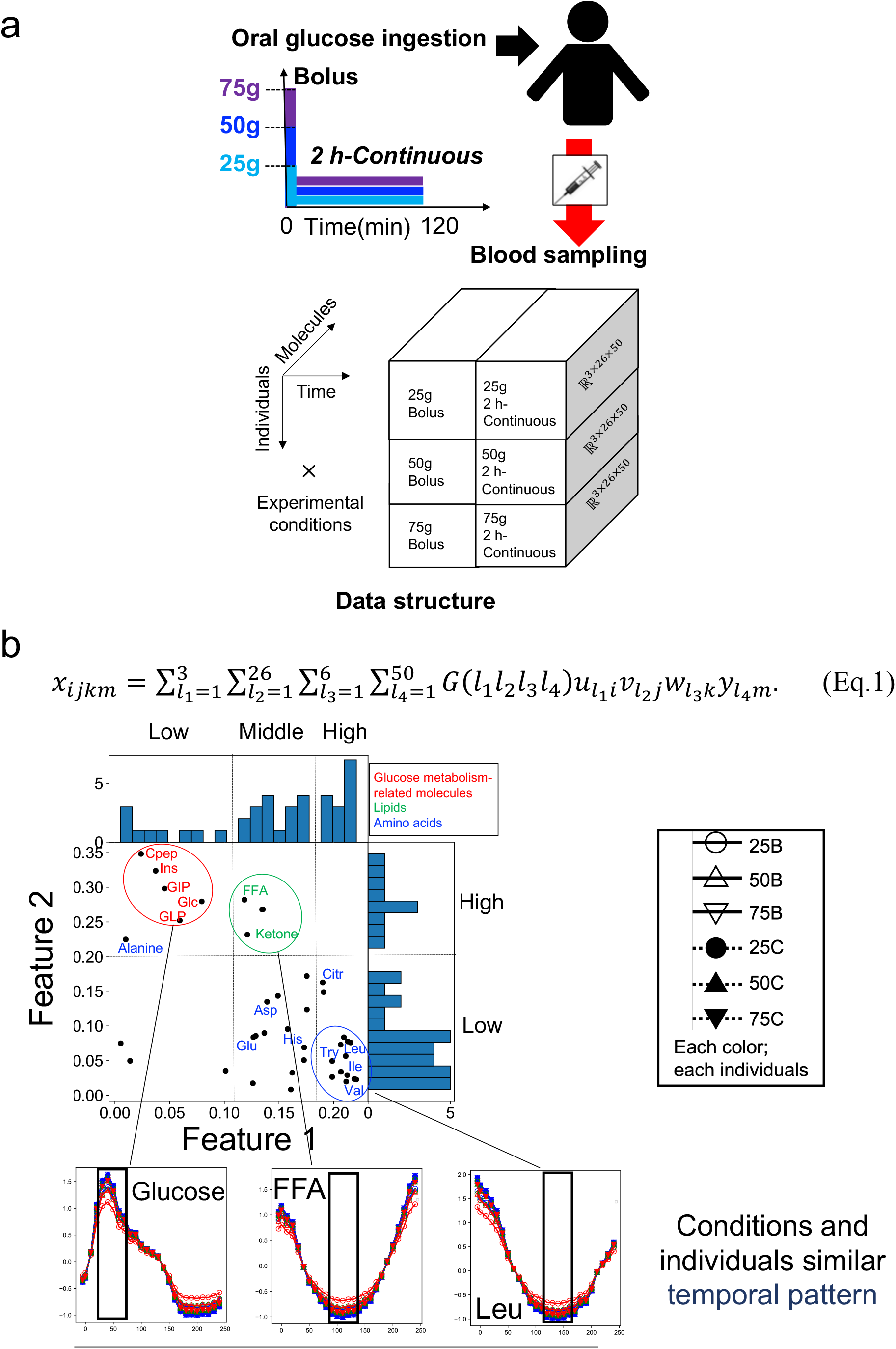
Data set for model cnstruction. **a** The experimental dataset for model selection. Three individuals orally ingested glucose with three doses 75, 50, and 25 g in two durations of bolus and 2 h continuous ingestion. The data structure has four axes: individual × time × experimental condition × molecule. The data represent the concentration changes at 26 time points (-5, 0, 10, 20, 30, 40, 50, 60, 70, 80, 90, 100, 110, 120, 130, 140, 150, 160, 170, 180, 190, 200, 210, 220, 230, 240 minutes) from 5 min before fasting to 240 min after glucose ingestion for 40 molecules in three healthy subjects, in six different experimental conditions. **b** Distribution of features extracted by tensor decomposition. Time series indicate similar temporal patterns among experimental conditions (‘Conditions similar’) and among individuals (‘Individuals similar’) for representative molecules^28^. The dashed lines show the values of Feature 1 divided into three, and Feature 2 divided into two, based on the shape of the distribution.The black lines in the time series indicate the time point of the peak. See our previous study for more details^28^. The color of each line indicates each condition. Solid lines indicate bolus ingestion, and dahed lines indicate 2 h continuous ingestion. Circles, triangles, and lower triangles indicate the three doses, 25, 50, and 75 g, respectively. Abbreviations for the representative molecules are as follows: Asp aspartic acid; Cit, citrulline; CRP, C-reactive peptide; FFA, free fatty acid; 3-OH, 3-hydroxybutyric acid; Ketone, Total ketone body; GIP, gastric inhibitory polypeptide (active); Glc, glucose; GLP, glucagon-like peptide-1; Glu, glutamic acid; His, histidine; Ile, isoleucine; Ins, insulin; Leu, leucine; Tyr, tyrosine; Val, valine. The label colors correspond to the metabolic group list (inset).

**Fig. S2.**
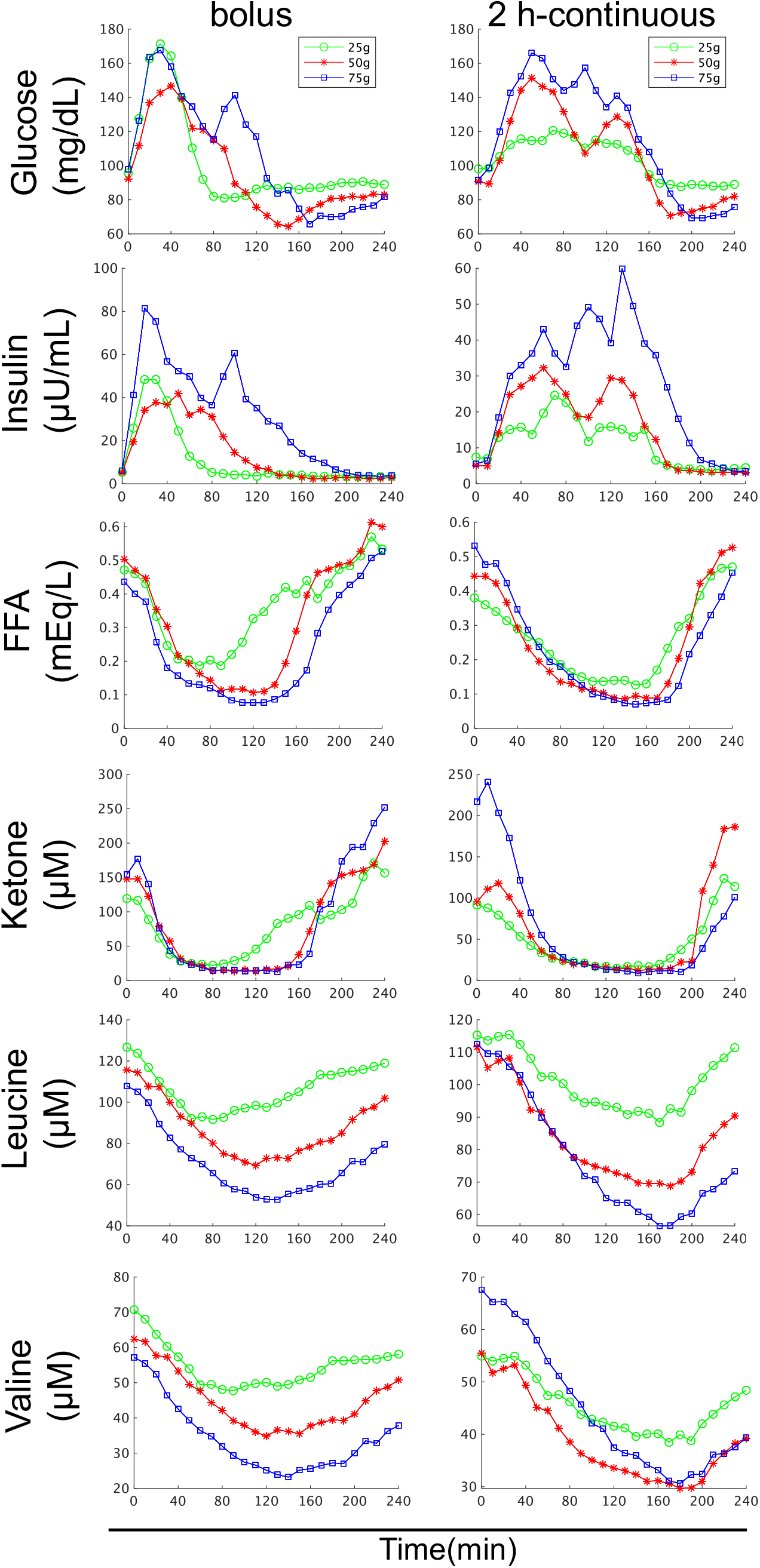
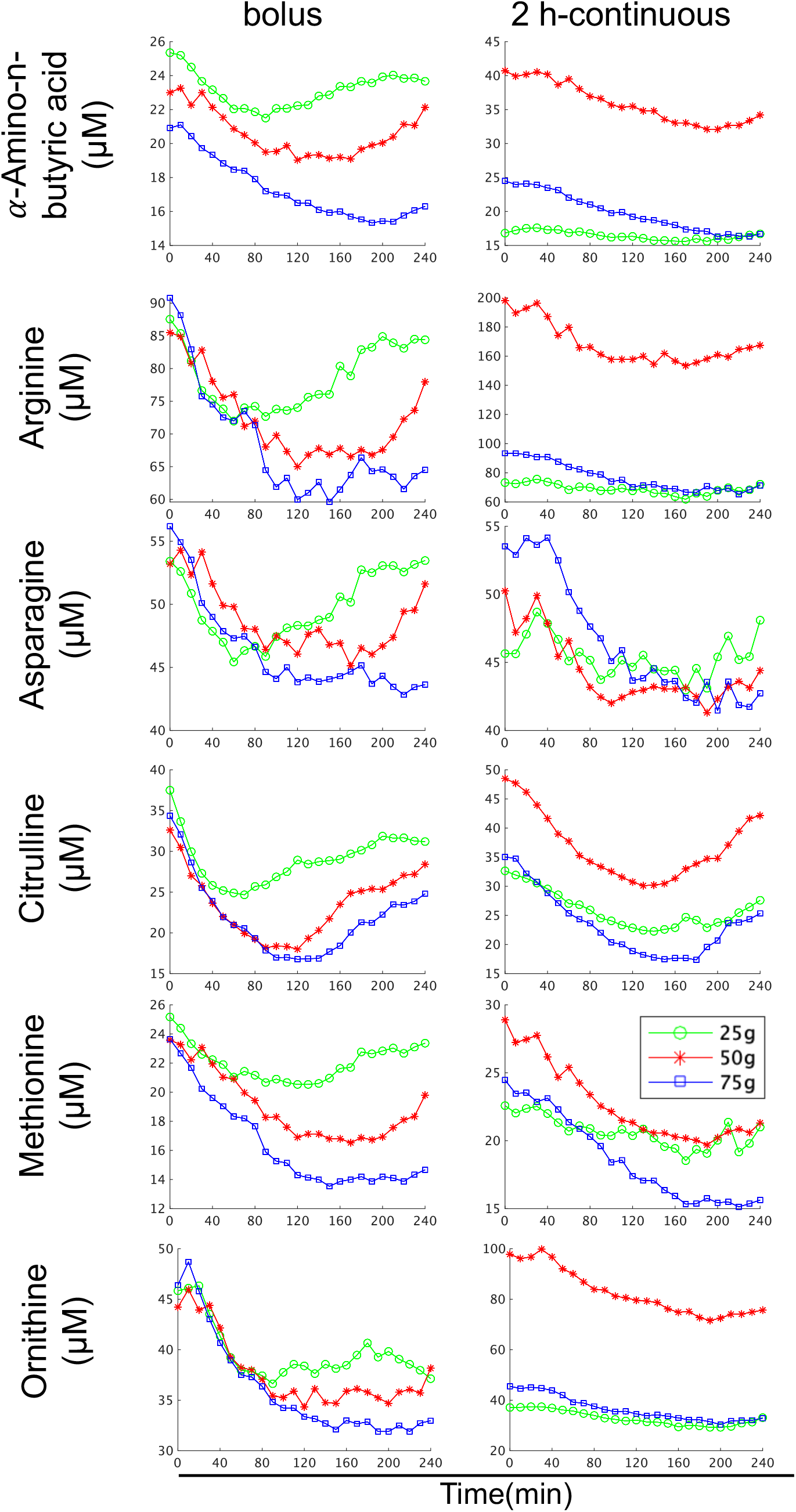
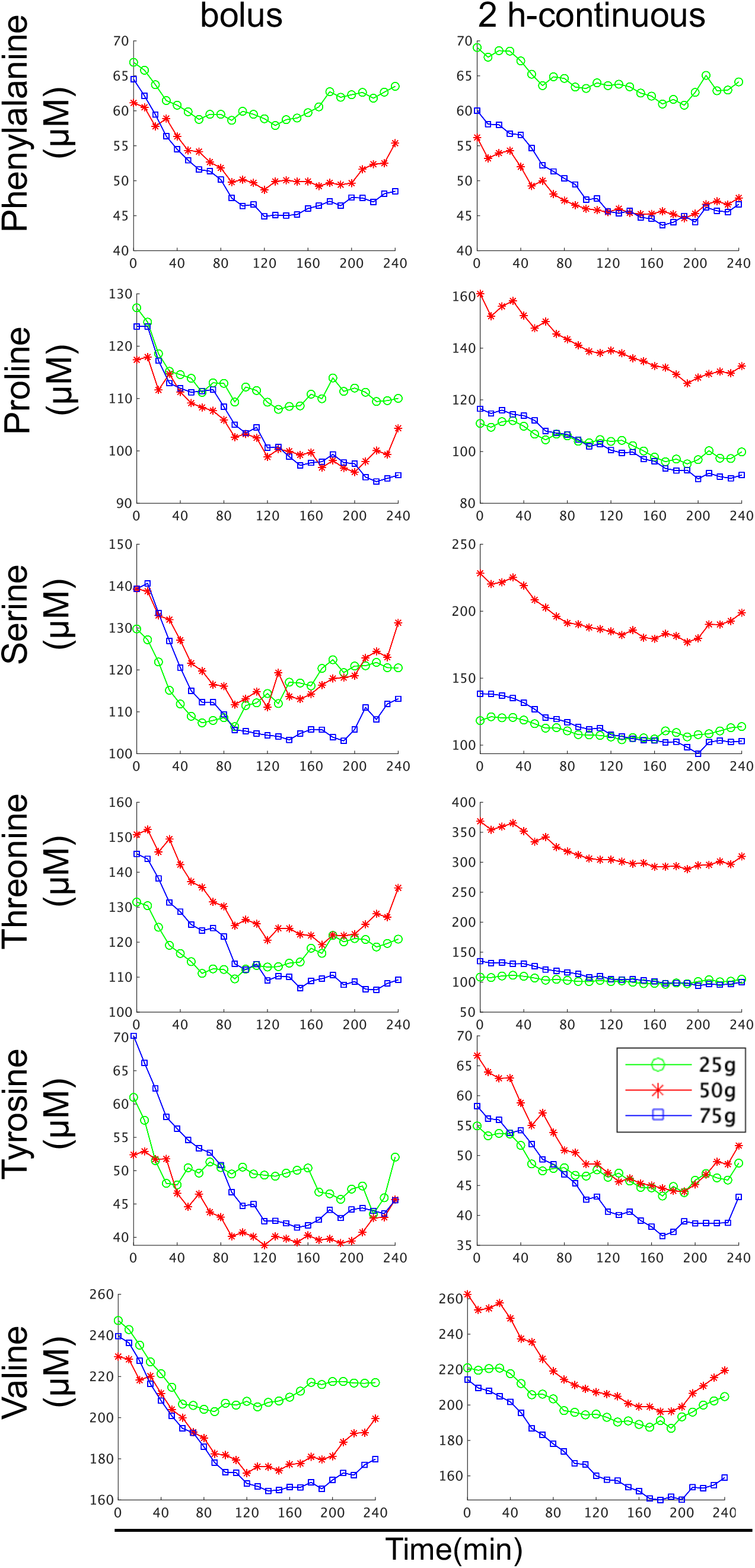
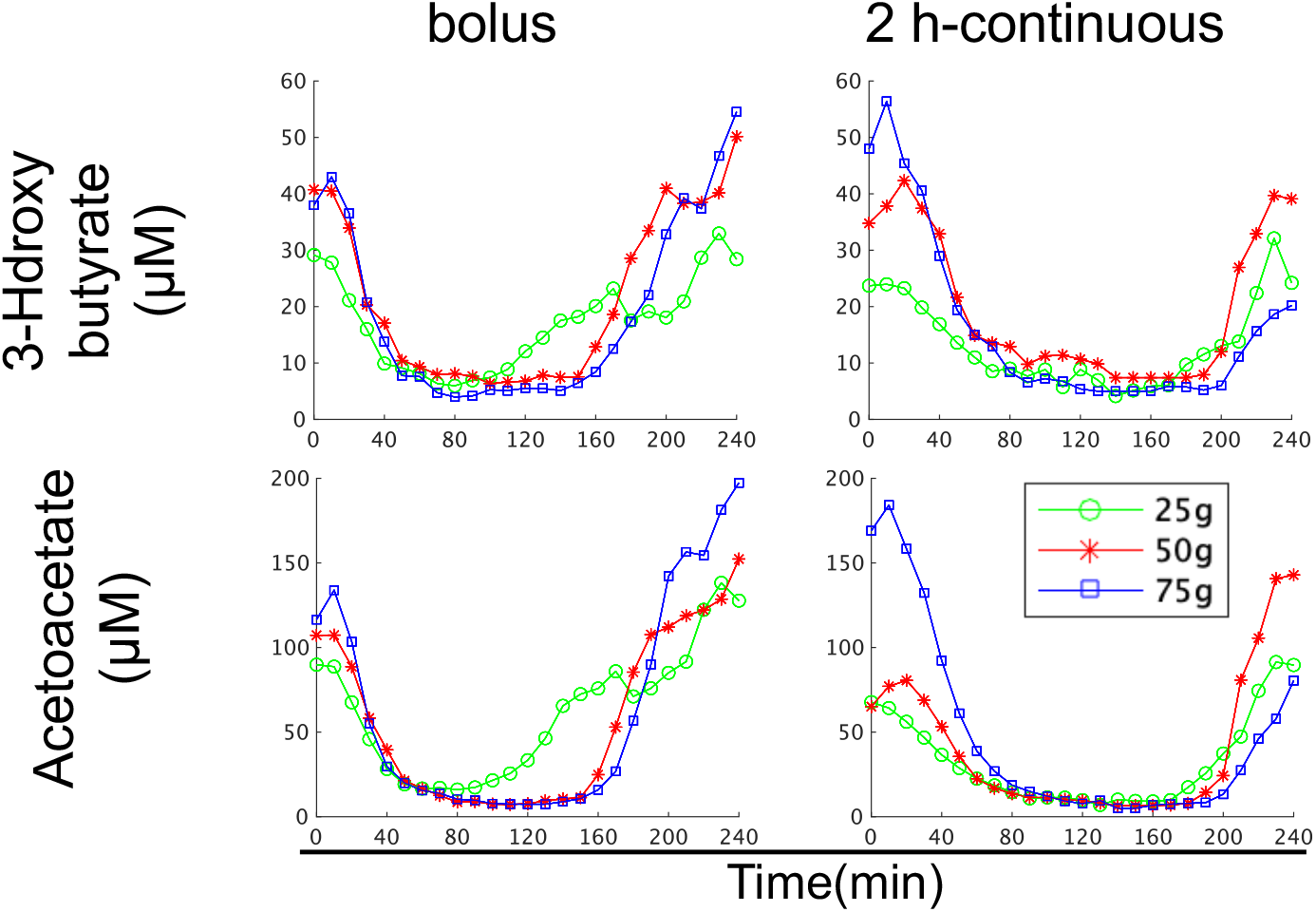
Time course data of blood insulin and blood metabolites in three individuals by glucose ingestion. The dose and ingestion pattern are indicated at the top. Green, red, and blue indicate three different doses: 25 g, 50 g, and 75 g, respectively.

**Fig. S3.**
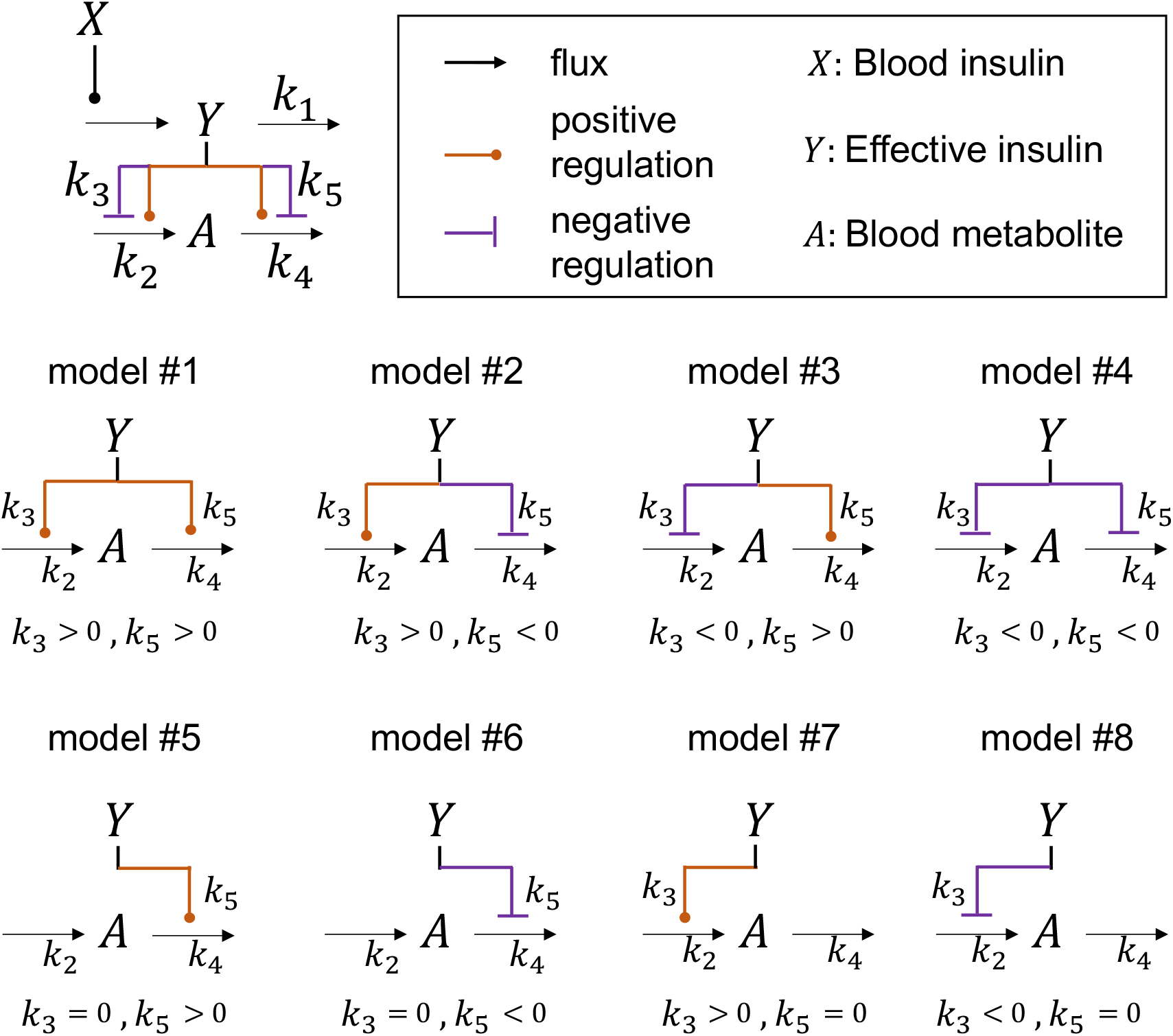
Model diagram.

**Fig. S4.**
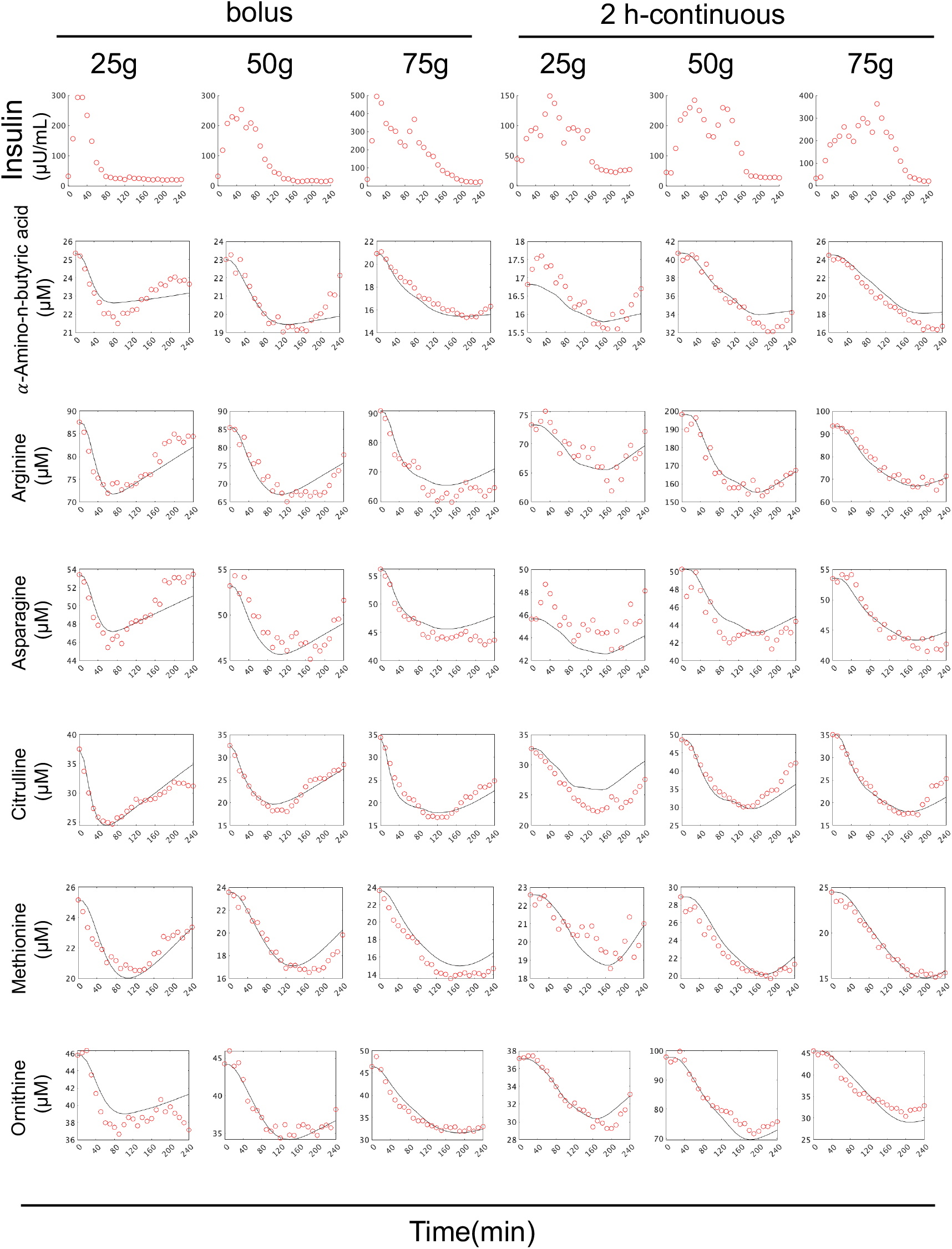
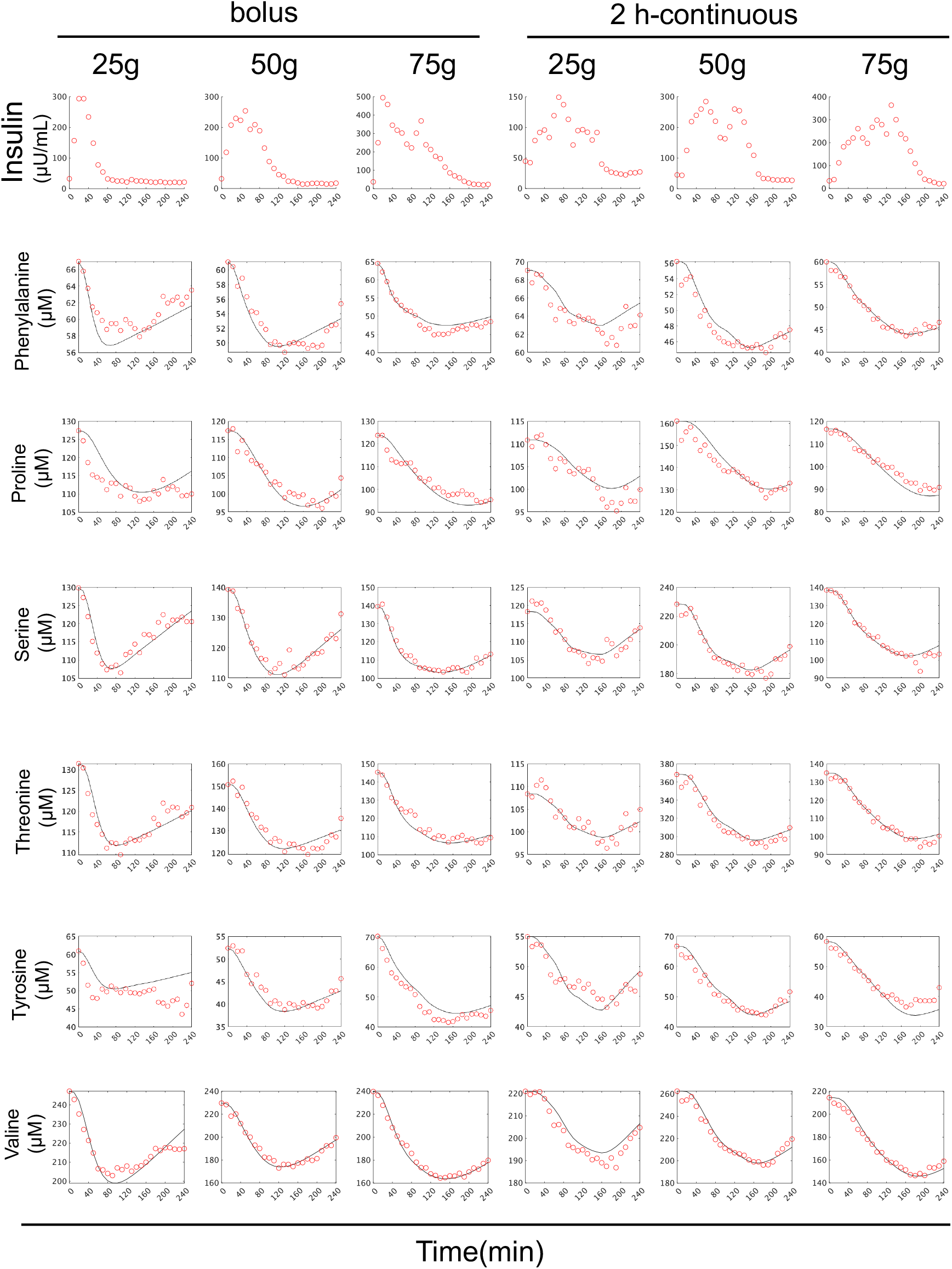
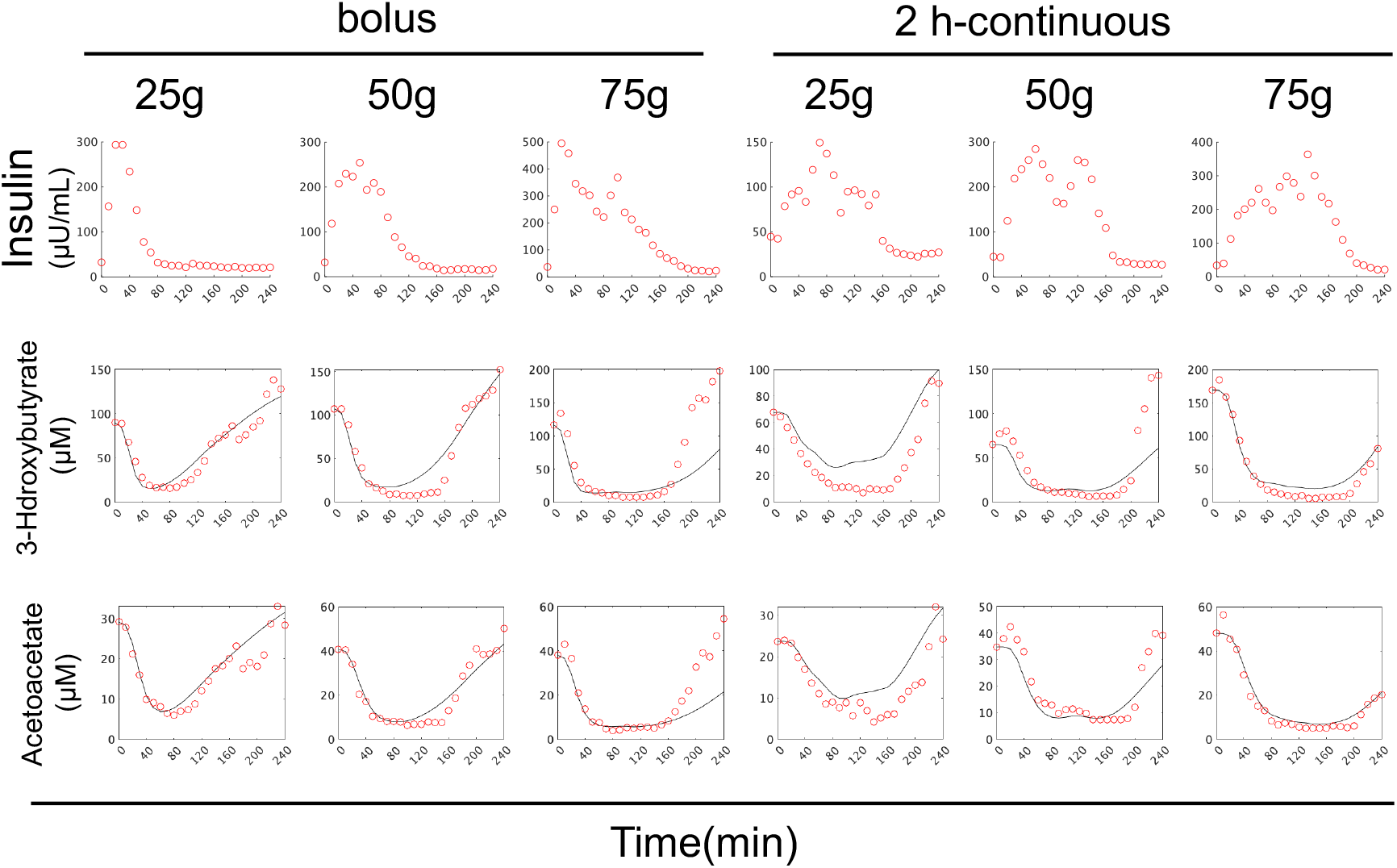
Time course data on the mean values of blood insulin and blood metabolites in three individuals by glucose ingestion. The doses and ingestion patterns are indicated at the top. The black lines indicate the temporal patterns of simulations, and the red circles indicate the time course data of experiments. The temporal patterns of the simulations were caluculated using the best fit model.

**Fig. S5.**
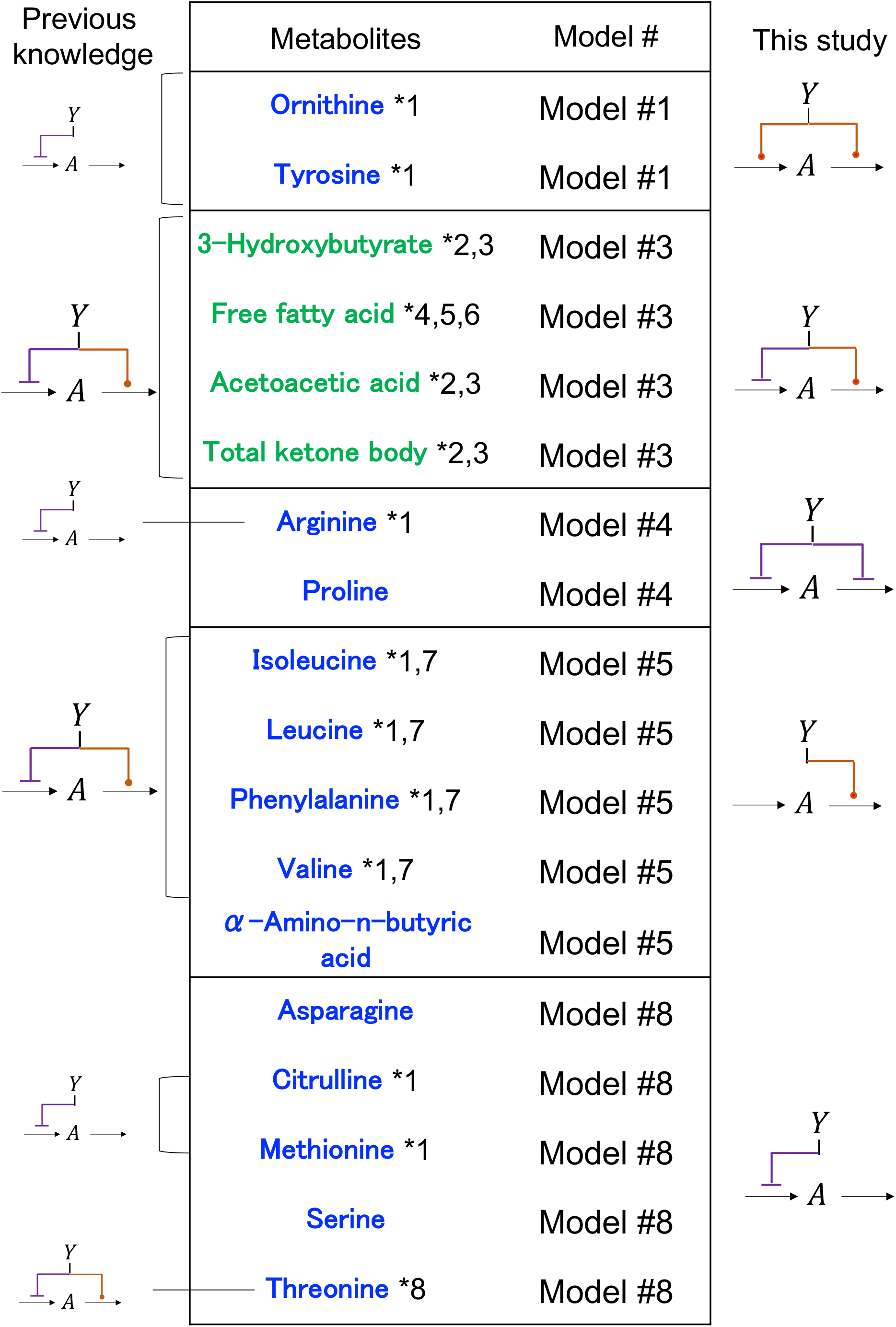
The model structures of of previous knowledge and the selected model of this study. Metabolites, numbers of selected models, model structures selected for this study and the regulatory structure of previous knowledge *1^4^, *2^23^, *3^26^, *4^40^, *5^41^, *6^5^, *7^3^, *8^19^.

**Fig. S6.**
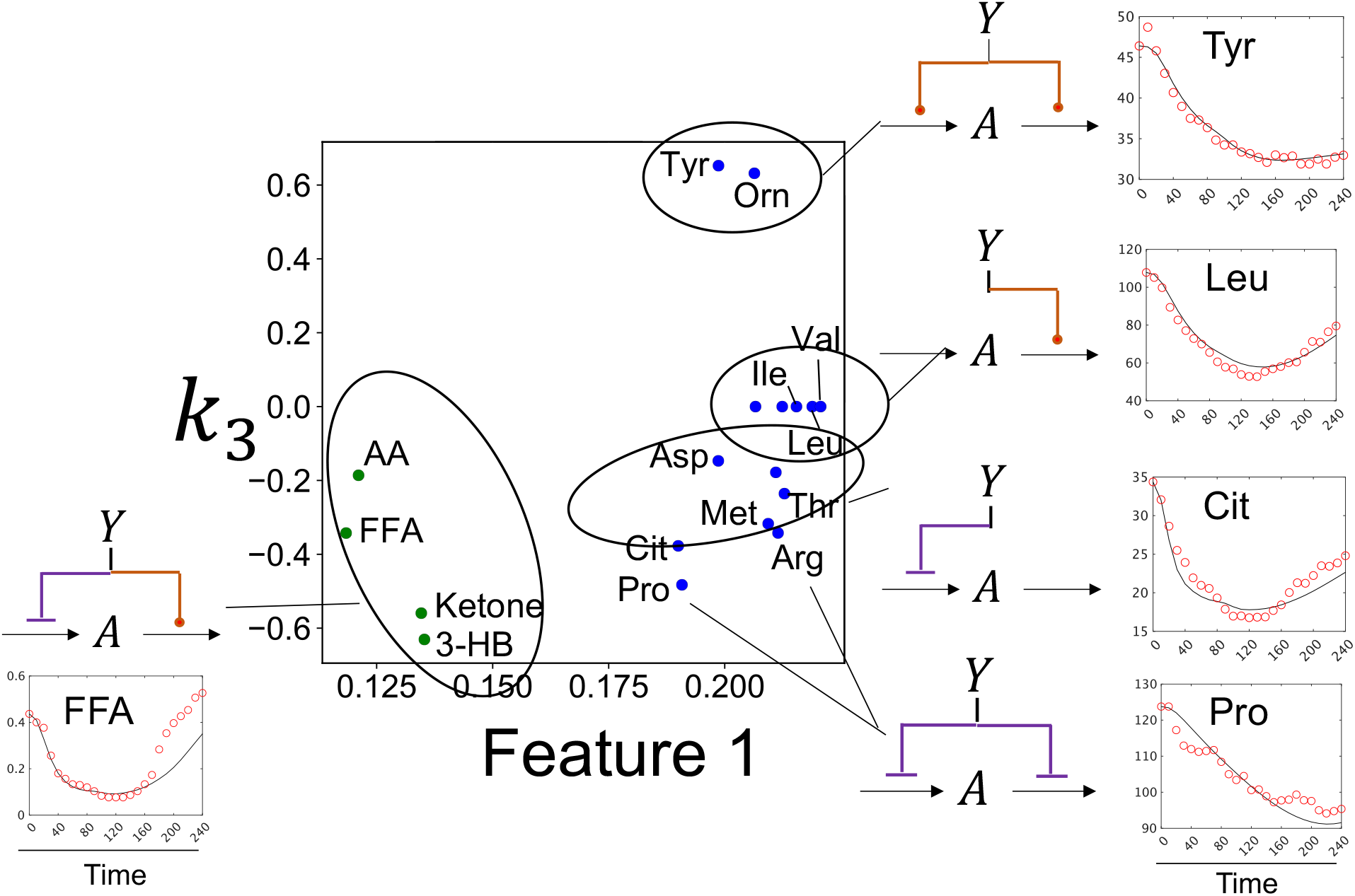
The features extracted by tensor decomposition. **a** Distribution of model parameters and Feature 1 (Fig. 3b). Circles and lines represent metabolites for which the same model structure was selected (Fig. 2). Time series of experimental and simulated values for a representative molecule for each selected model are shown (75 g bolus).

**Fig. S7.**
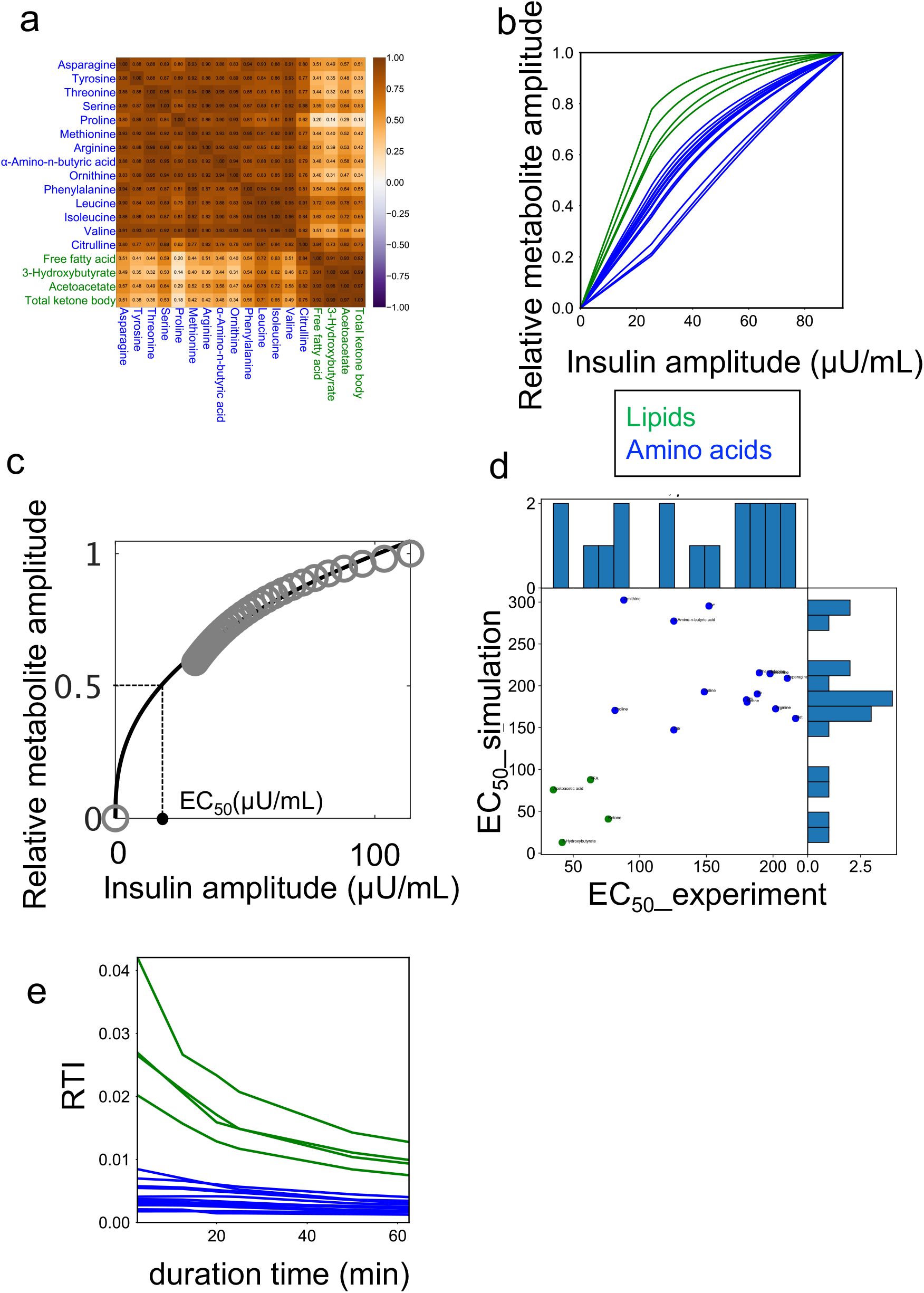
The regulation of the amplitude of amino acids and lipids by amplitude of insulin. **a** Heat map showing the temporal pattern similarity among metabolites among all metabolites(see Methods). The color of the letters indicates metabolic group (blue: amino acids, green: lipids) **b** The peaks of all metabolite against the amplitude concentration of insulin in the simulation. The color of the line indicates the metabolic group. **c** The definition of EC_50_ against the amplitude concentration of insulin. **d** Distributions of experimental EC_50_ value and simulated EC_50_ value of metabolite. As with the simulated EC_50_ value, nonlinear fitting was performed to calculate the experimental EC_50_ value using the maximum value of insulin in the time series and the log2 Fold change relative to the metabolite fasting value. **e** RTIs for the metabolite against the duration time(inverse of rate of response) of insulin.

**Fig. S8.**
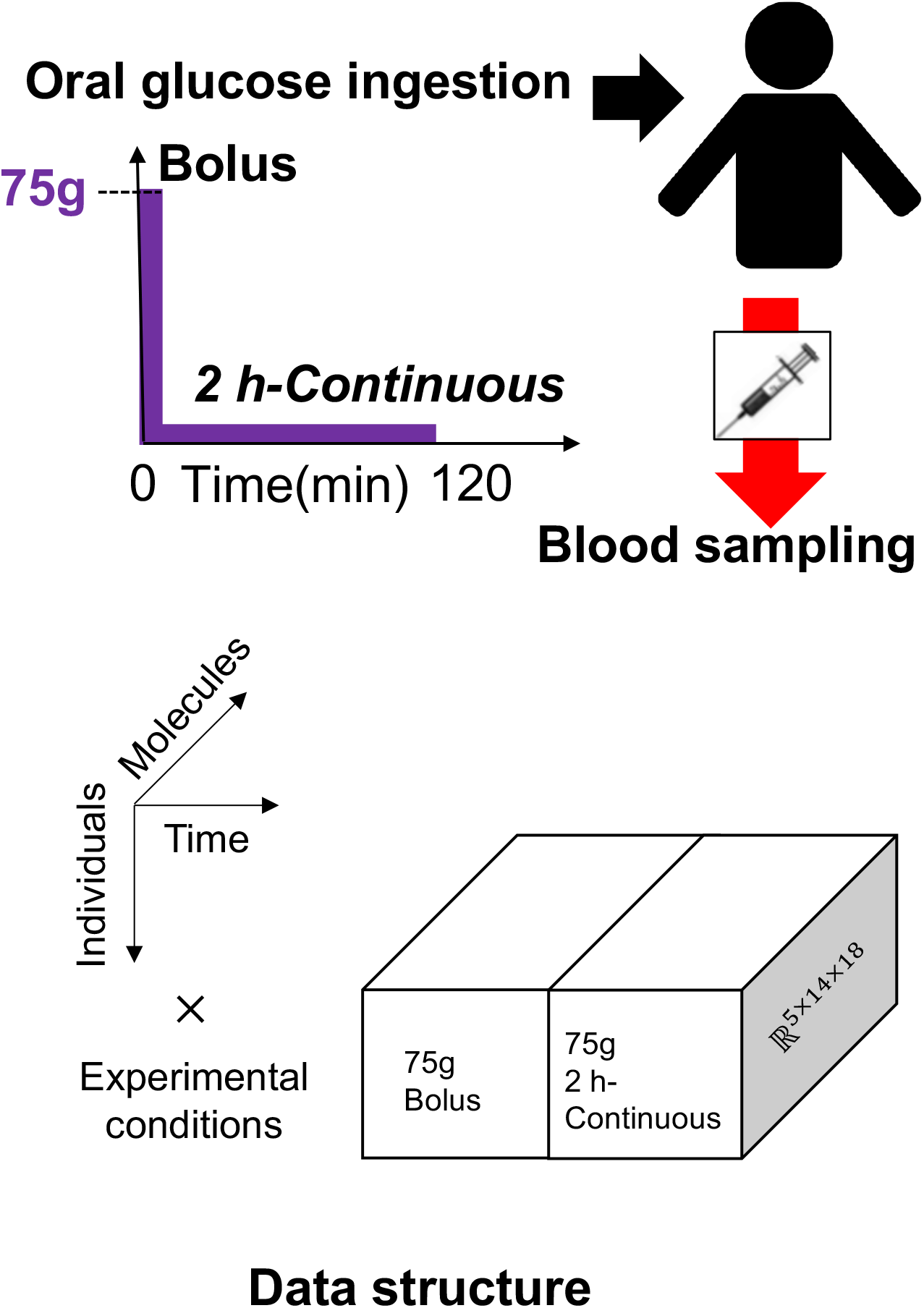
Data set for model validation. Experimental dataset for validation. Five individuals orally ingested glucose with three doses 75g in two durations of bolus and 2 h continuous ingestion. The data structure has four axes: individual × time × experimental condition × molecule. The data represent the concentration changes at 26 time points (-5, 0, 10, 20, 30, 40, 50, 60, 70, 80, 90, 100, 110, 120, 130, 140, 150, 160, 170, 180, 190, 200, 210, 220, 230, 240 minutes) from 5 min before fasting to 240 min after glucose ingestion for 20 molecules in fivehealthy subjects, in six different experimental conditions.

**Fig. S9.**
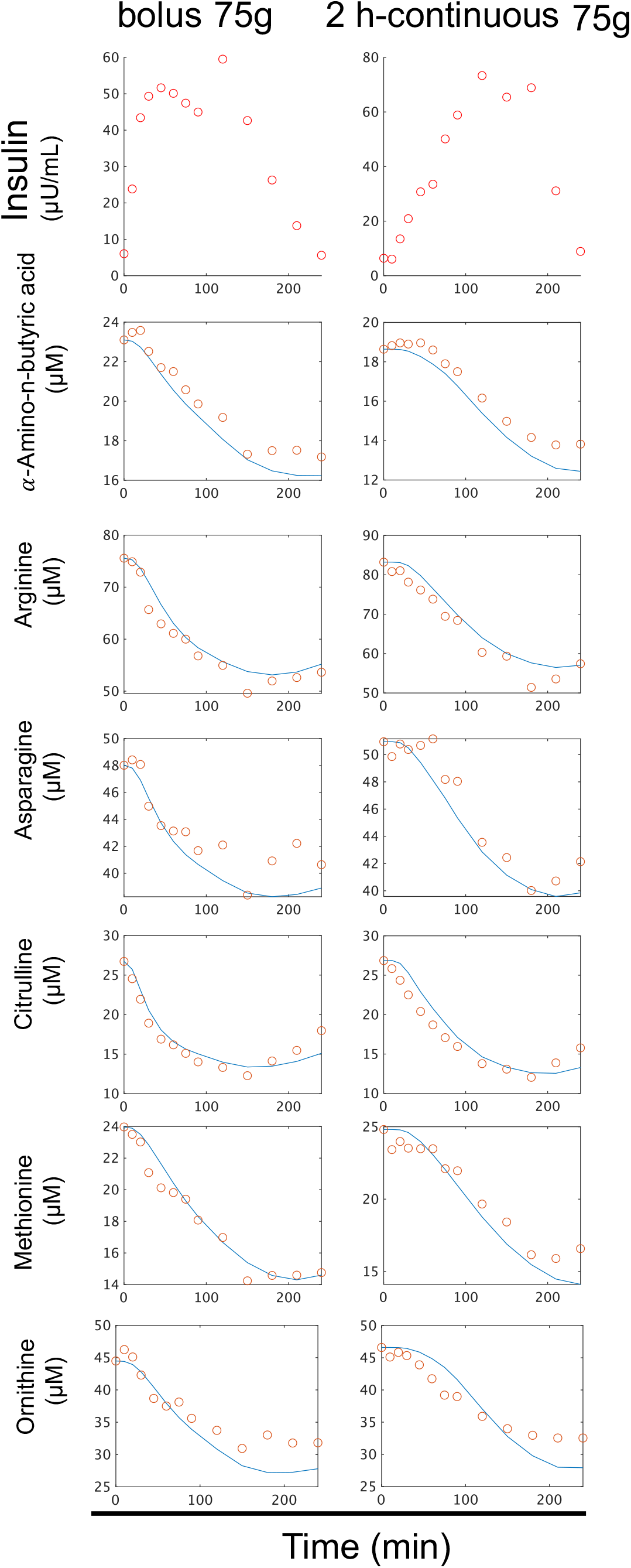
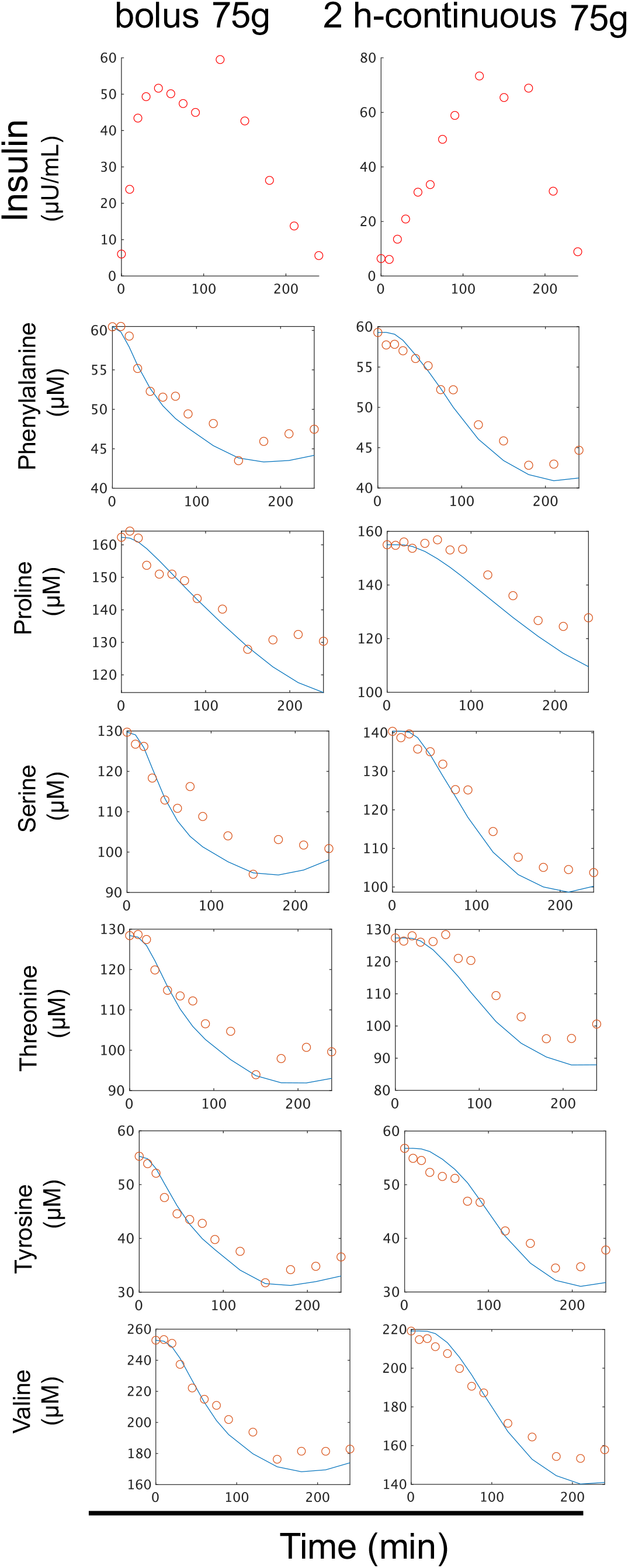
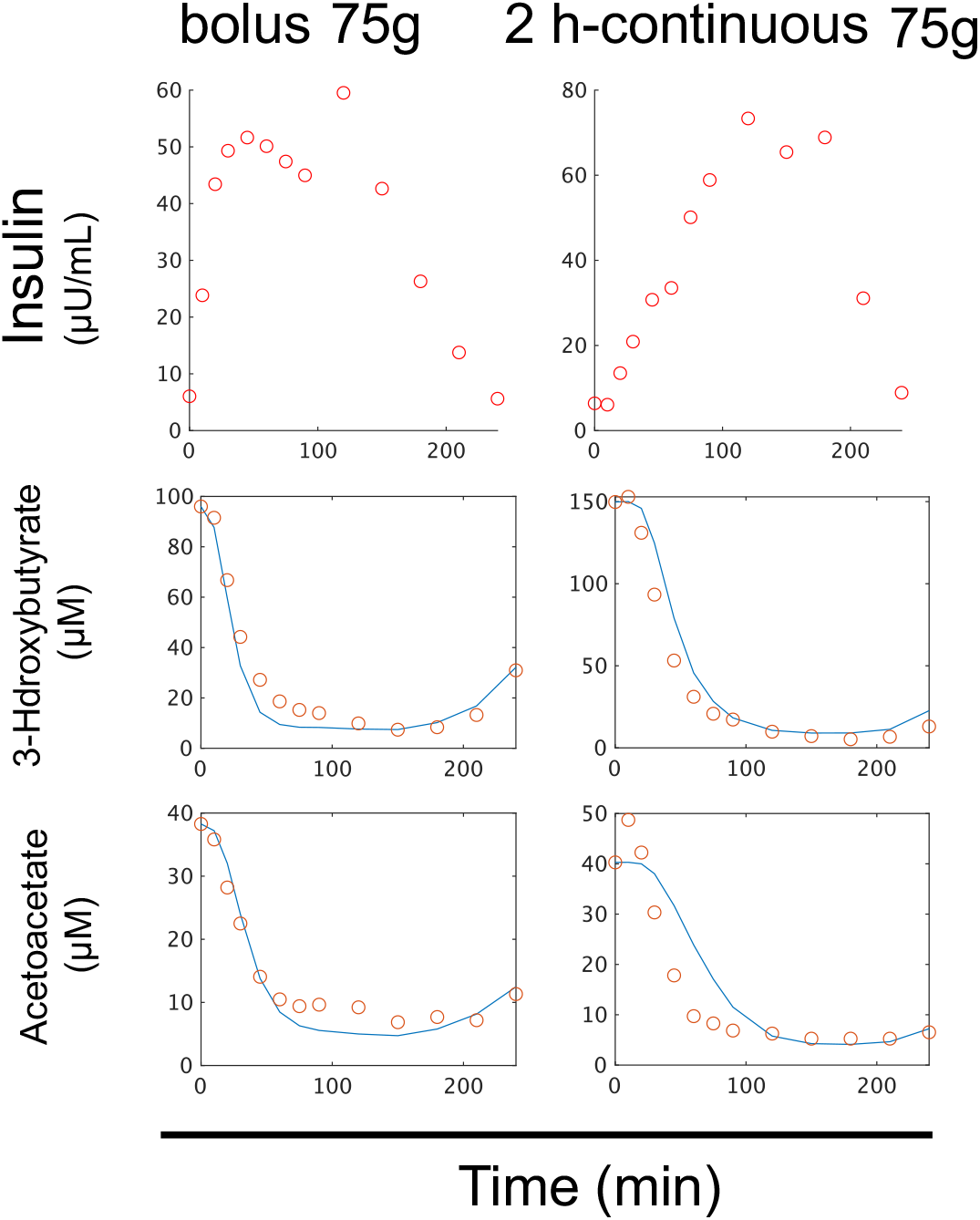
Time course data on the mean values of blood insulin and blood metabolites in five individuals by glucose ingestion for model validation. The dose and ingestion pattern are indicated at the top. The bluelines indicate the temporal patterns of simulations, and the red circles indicate the time course data of experiments.

**Fig. S10.**
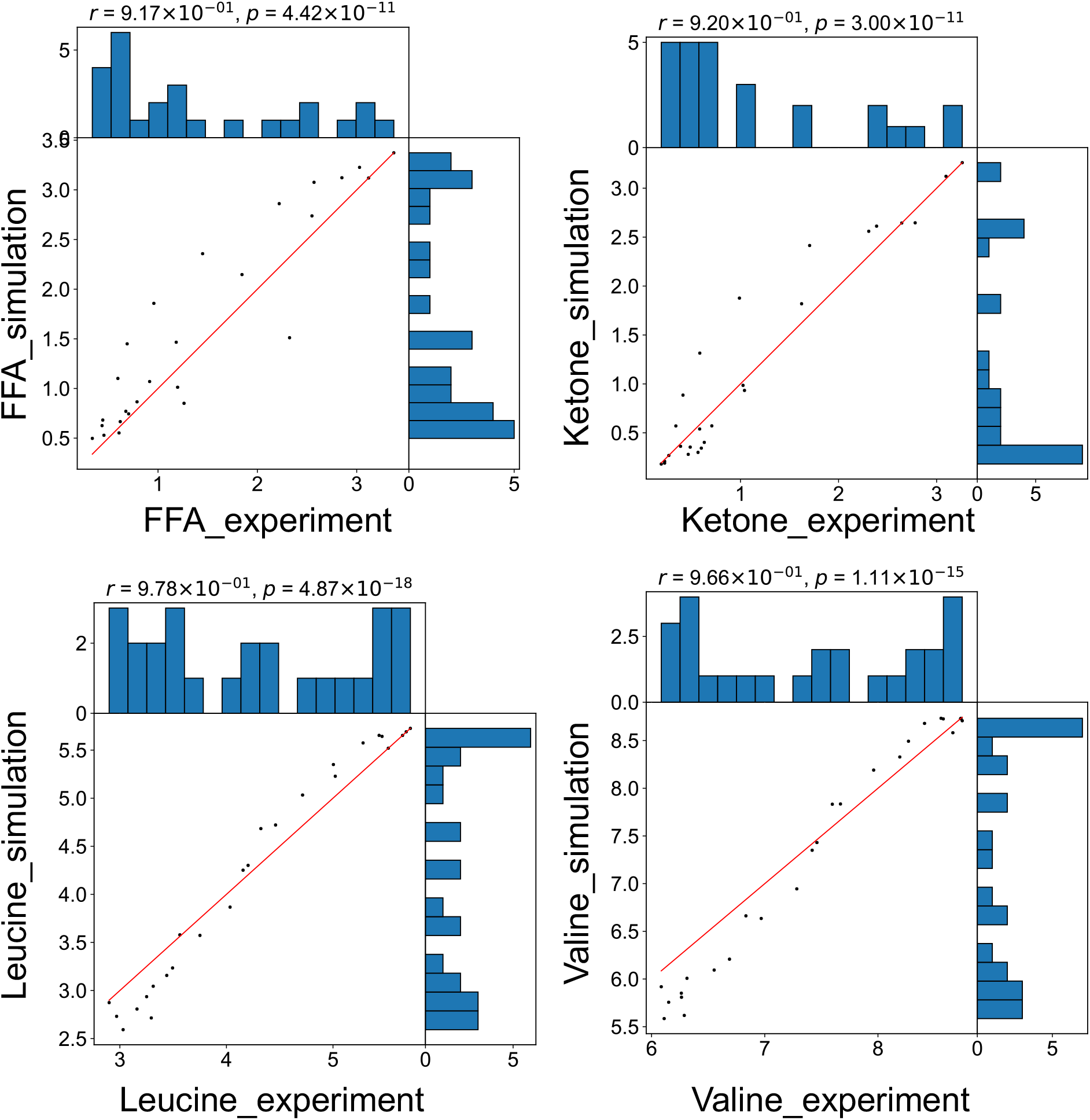
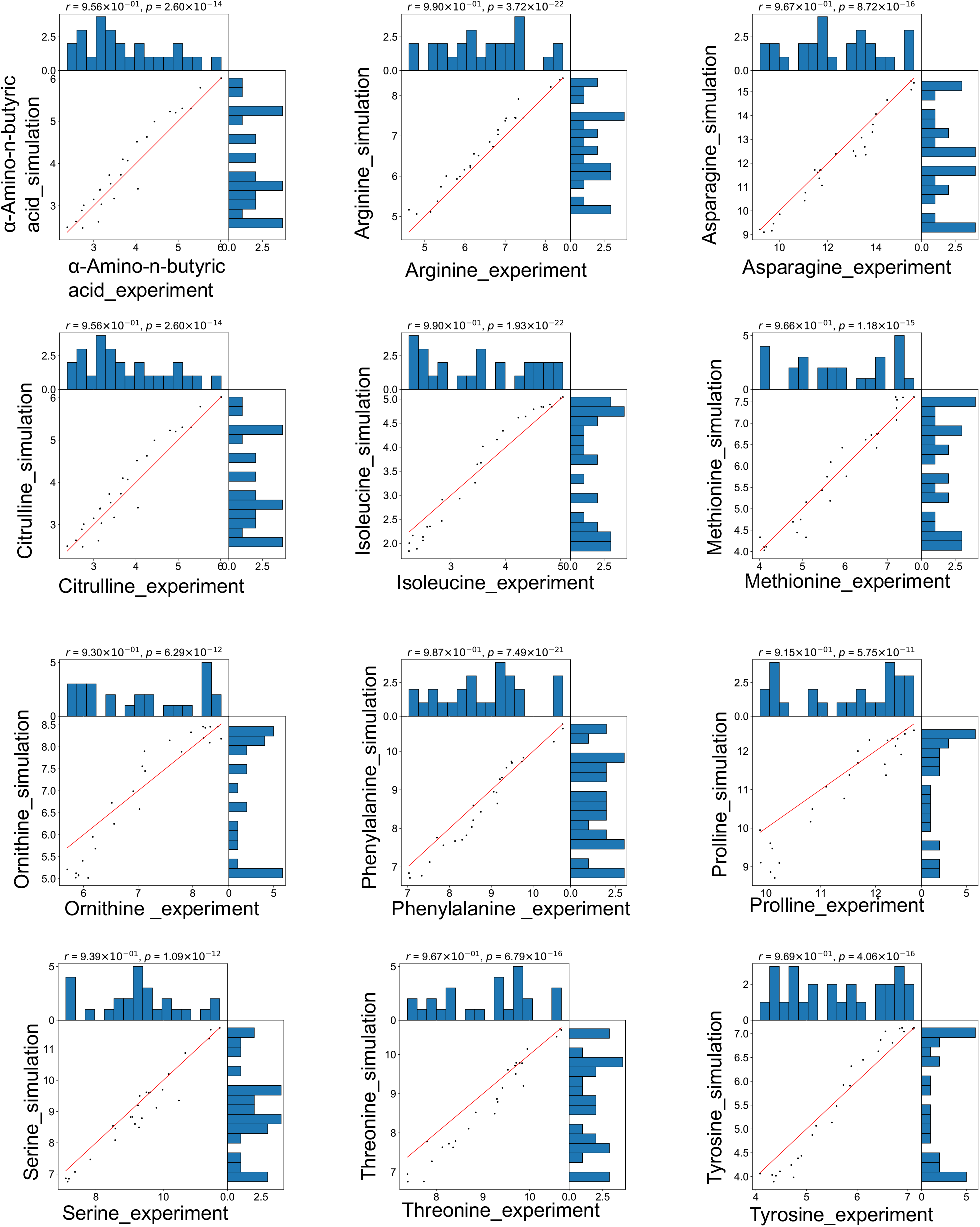
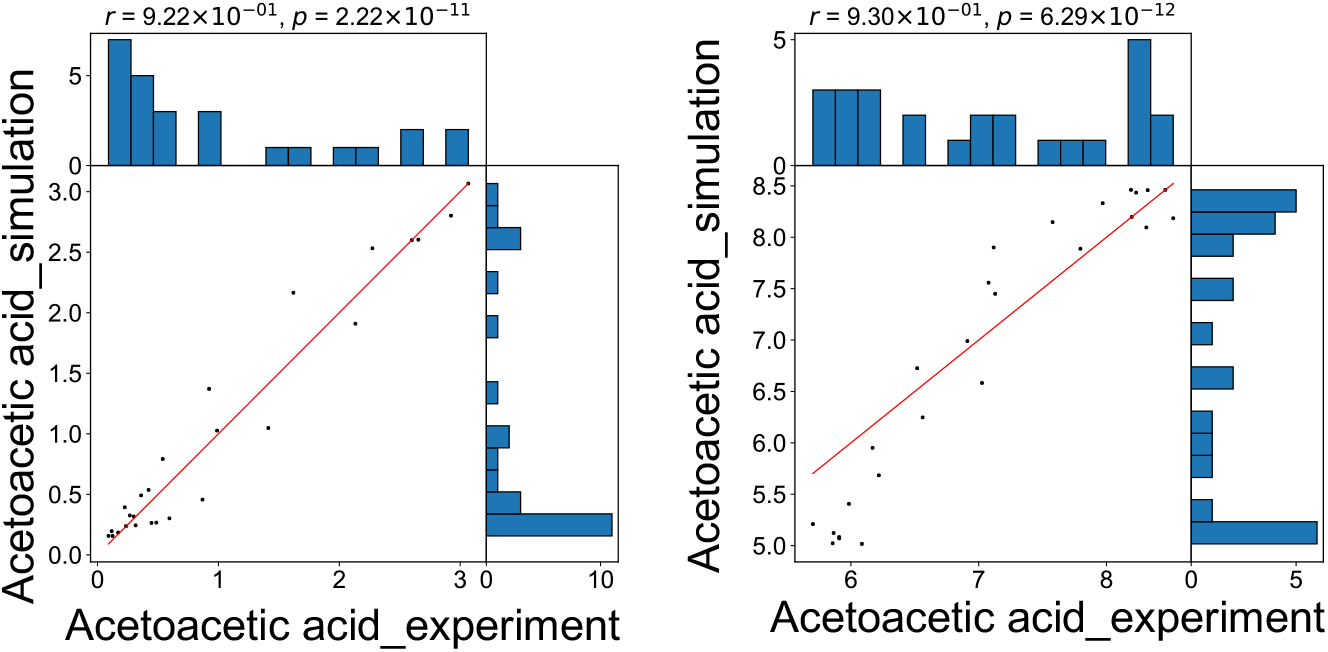
Distributions of experimental value and simulated value of metabolite. In the upper right, *r* and *p* indicate the correlation coefficient and *p* value, respectively. The red line indicates y=x.

**Fig. S11.**
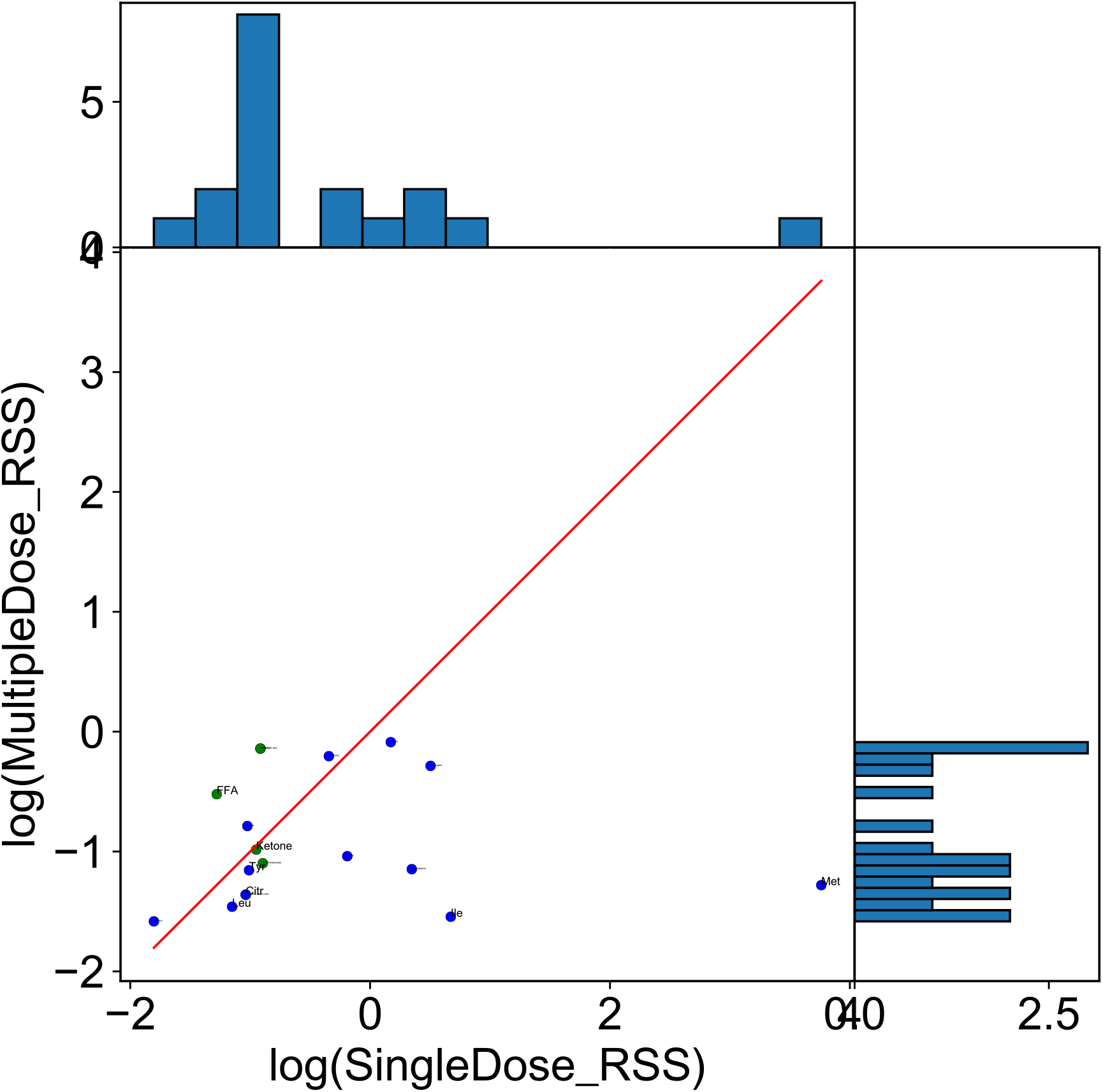
Influence of learning data on prediction. The rss between simulations with parameters estimated from experimental data for 75 g, 50 g, and 25 g bolus ingestion and 2 h-continuous ingestion (all experiments, y axis) and 75 g bolus ingestion only (75 g bolus only, x axis), and experimental values for all metabolites.The red line indicates y=x. The dot’s colors correspond to the metabolic group (blue: amino acids, green: lipids)

**Fig. S12.**
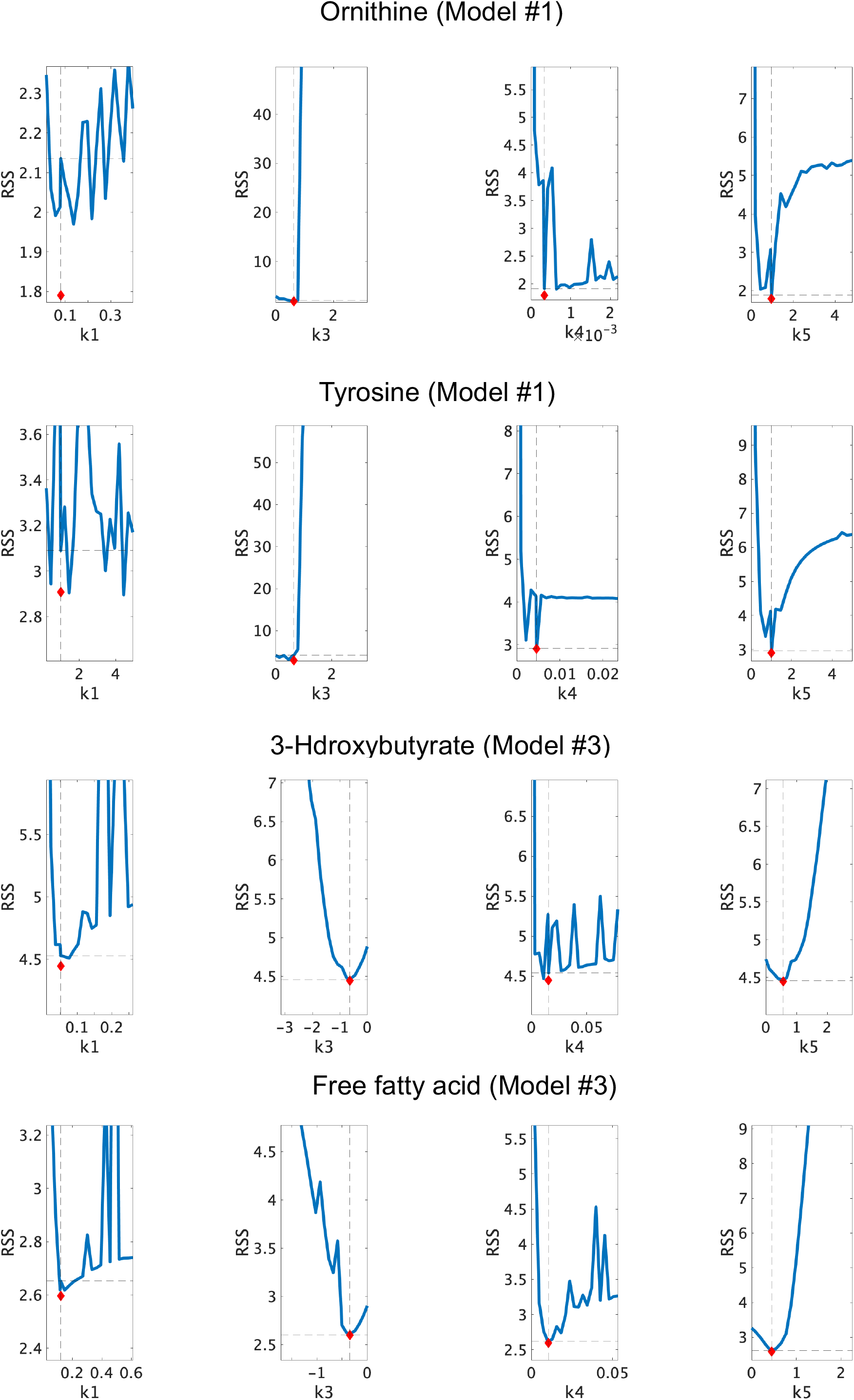
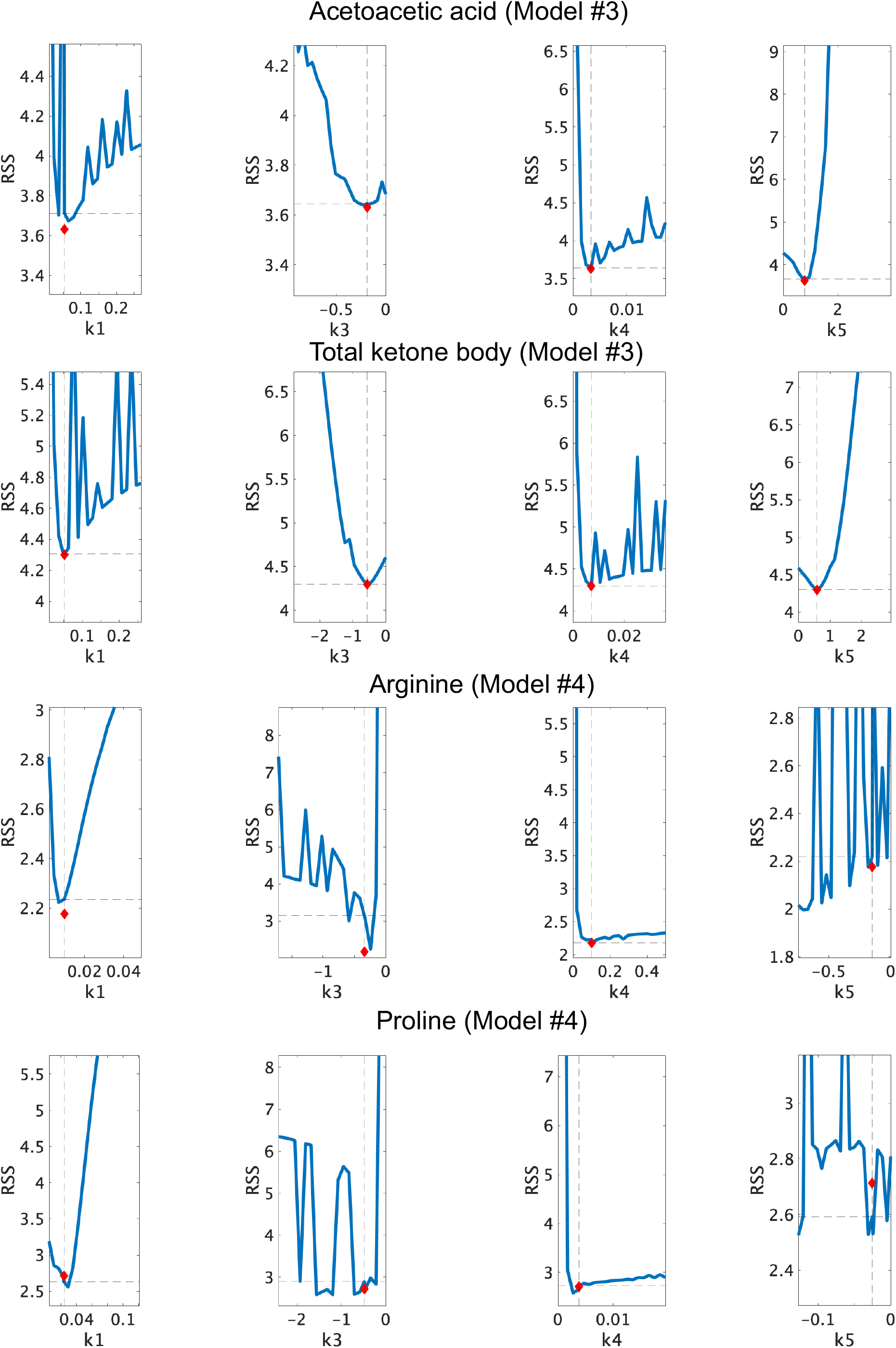
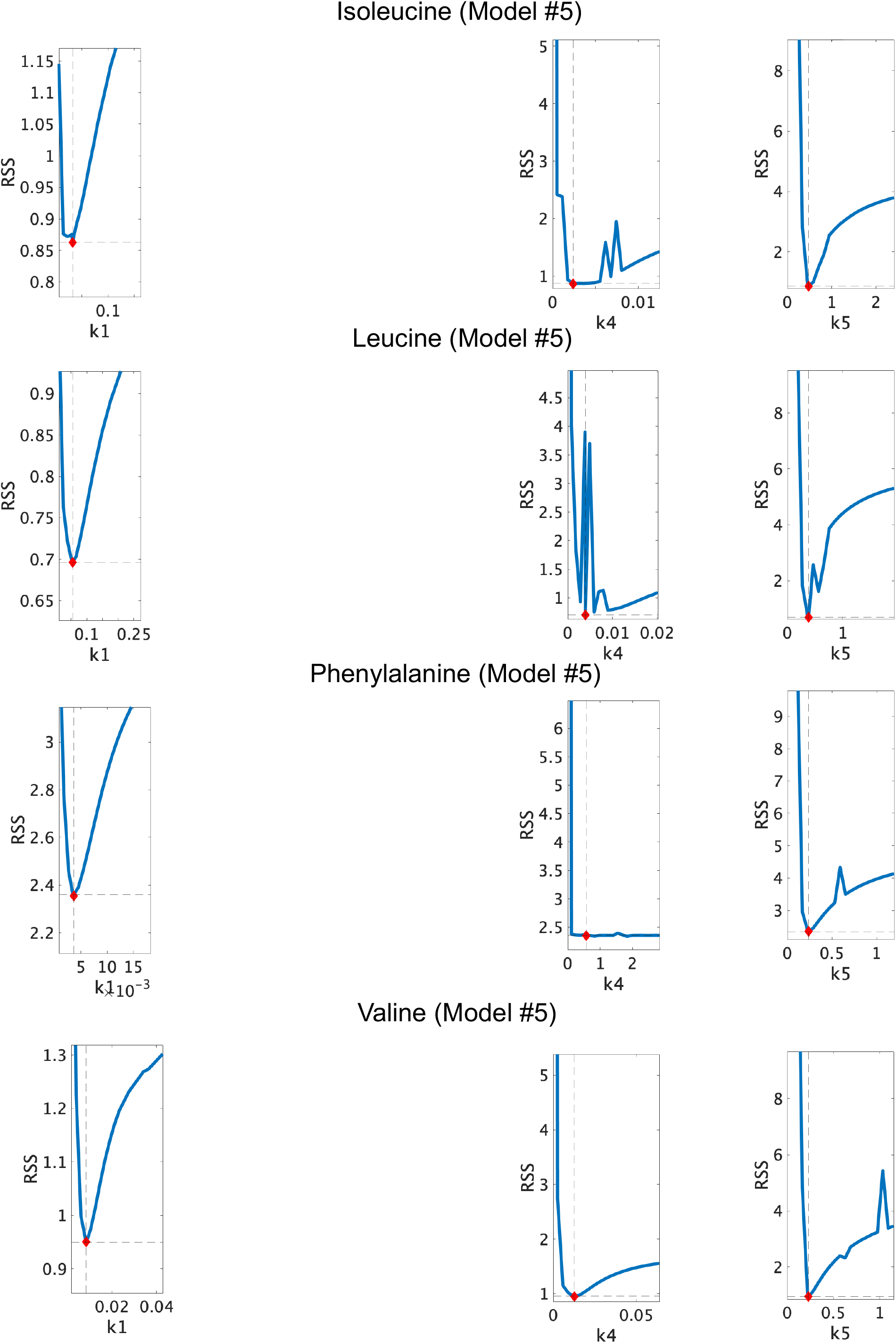
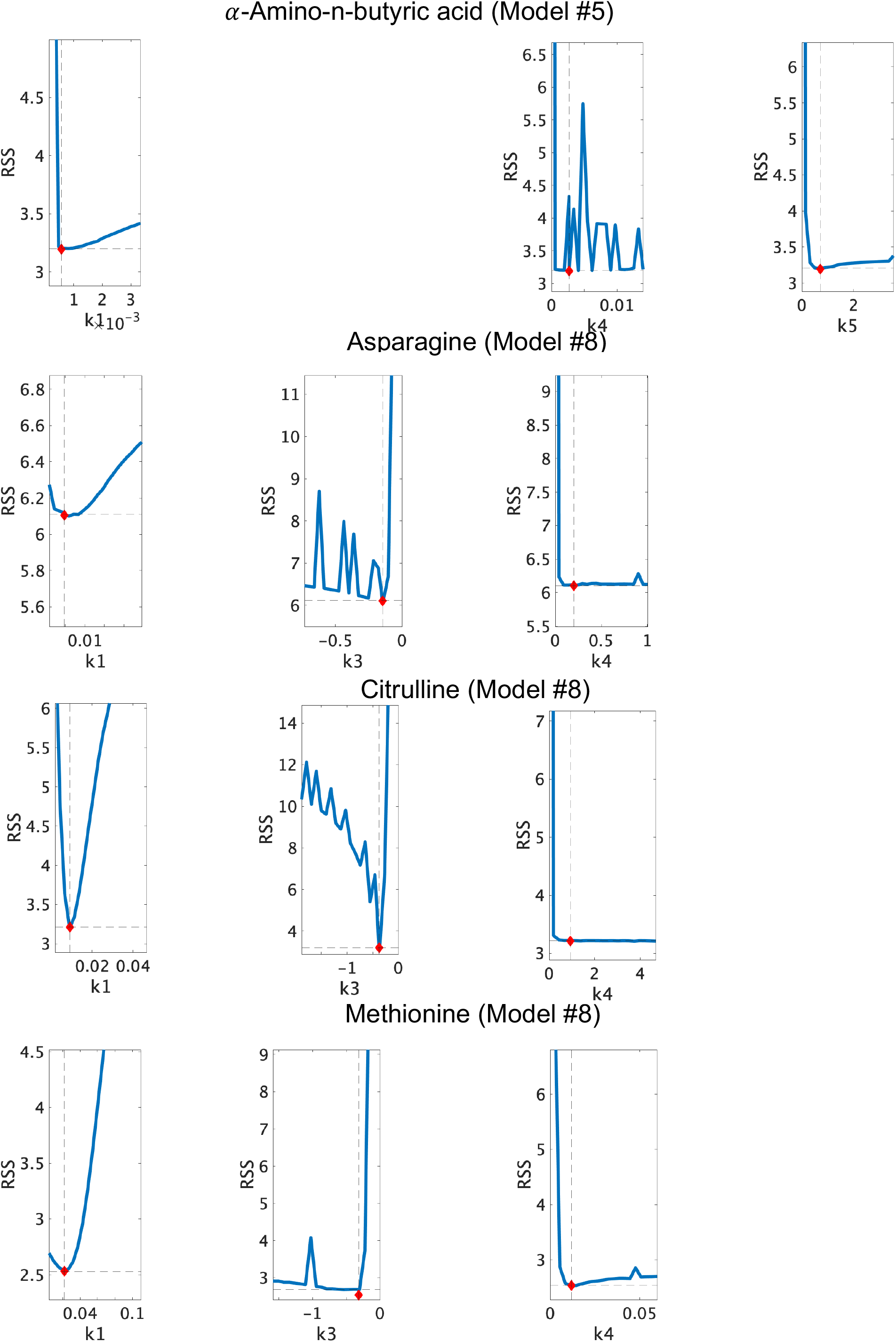
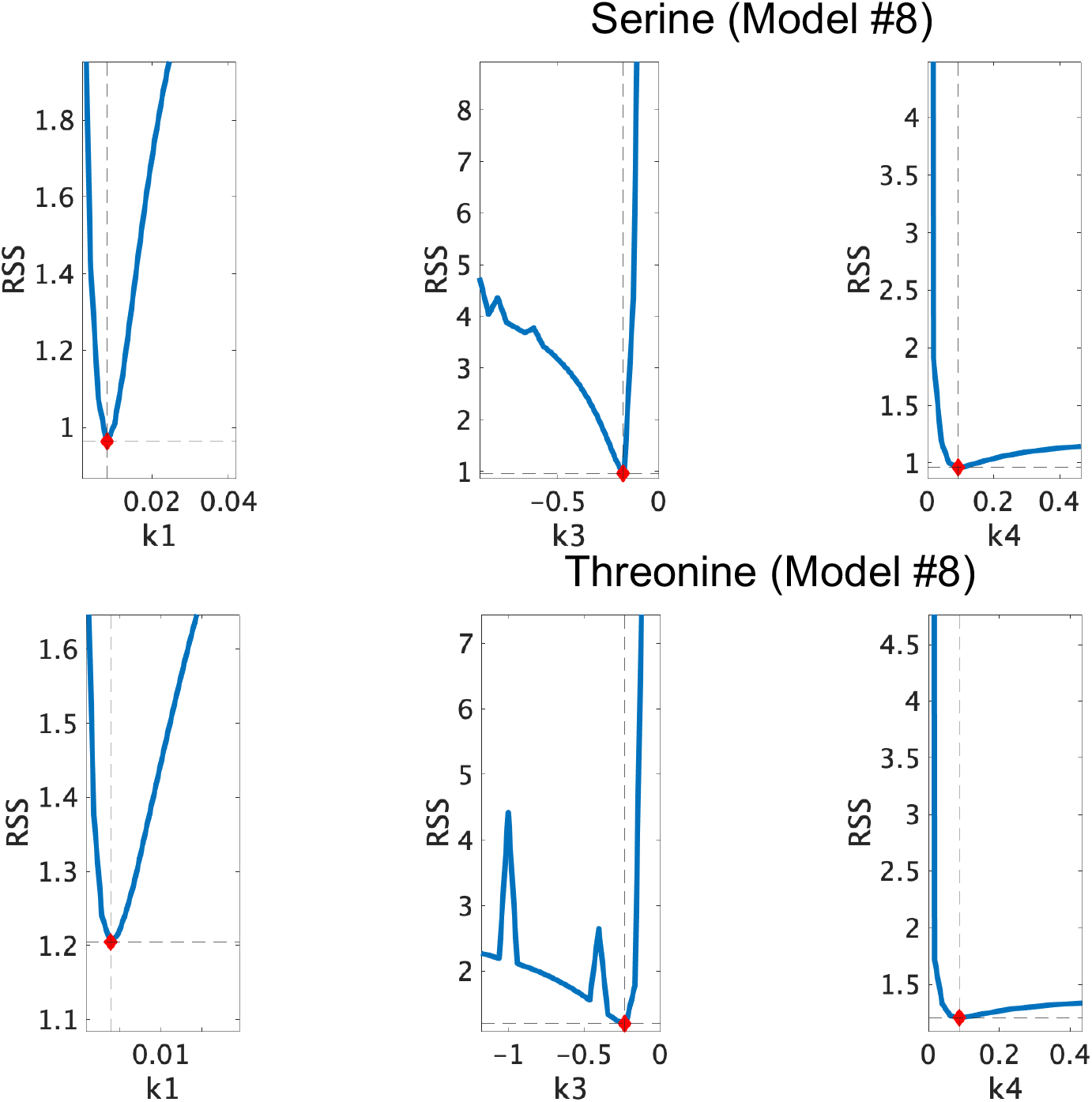
Identifiability of the parameters in the model structures selected for each metabolite. The red star indicates the RSS of the model fitted using the optimal parameter values estimated from the data, and the blue line indicates to the RSS when other parameter values are re-estimated after iterative adjustment of the parameter values. The dashed line represents the RSS when re-estimated using the same values as the optimal parameter values. Parentheses denotes the selected model.

## Notes

### Competing Interest Statement

The authors have declared no competing interest.

